# Substantial Deceleration of Adaptation of HIV-1 Within 1,500 Generations in an Experimental Evolution: A Genomic Perspective

**DOI:** 10.64898/2026.02.04.700995

**Authors:** Ali Movasati, Christine Leemann, Kathrin Neumann, Rongfeng Chen, Lygeri Sakellaridi, Karin J. Metzner, Roland R. Regoes

## Abstract

Numerous experimental evolution studies have suggested that adaptation rate of microbial populations evolving in stable environments decline over time. Despite the generality of this phenomenon across different domains of life, the timing and magnitude of decline in adaptation can vary greatly based on the idiosyncrasies of the biological system. To investigate the characteristics of adaptation deceleration in a fast-evolving virus, we propagated HIV-1 in two human T-cell lines (MT-2 and MT-4) for approximately 4.8 years and tracked its genome evolution through next-generation sequencing. The curated sequencing data covering the whole-genome can be accessed and explored via LTEEviz, an interactive web application. Time-resolved sequencing data uncovered that despite constant fixation rate of 0.085 (MT-2) and 0.042 (MT-4) mutations per generation, the fixation kinetics of adaptive mutations changed considerably over time. The rate of fixation of adaptive parallel mutations decreased by 44% per 300 generations, while their conferred fitness gain decreased by 27% (MT-2) and 18% (MT-4) per every added adaptive mutation in their genetic background. The early and substantial deceleration of adaptation in our HIV-1 populations can, at least in part, be explained by diminishing gains of adaptive mutations. Furthermore, we identified genomic patterns consistent with a hard selective sweep that occurred in one population later in the experiment. Together, our results confirm that HIV-1 genomic evolution is characterized by a swift and substantial deceleration of adaptation, while also revealing that episodes of positive selection can occur beyond the initial adaptive phase.

## Introduction

Decades of evolutionary research, spanning from theoretical models to empirical studies, have indicated that the fitness of evolving microbial populations in a stable environment effectively reaches a plateau^1–3^. Despite the predictable decline in the rate of adaptation in constant environments, the trajectories and timescales by which populations approach the fitness optimum can vary greatly^3–6^. Stochasticity in evolutionary processes permits populations to take unique paths through genotype space^7^, while the rate of adaptation is influenced by several factors such as availability of genetic variation^8^, the demographic properties of populations^9^, and the topology of the fitness landscape^10^. Thus, a full understanding of the mode and tempo of adaptation and its determinants requires examination of adaptive dynamics in a wide range of organisms.

One approach to studying adaptive dynamics of evolving populations is to probe patterns of polymorphism and divergence at the nucleotide level^11^. These patterns are shaped by the interplay of multiple evolutionary forces, including selection as the propellant of adaptation^12^. When populations approach an adaptive peak, the influence of positive selection on genome evolution wanes^13^, while other background evolutionary forces continue to operate unabated^14,15^. Hence, identifying and gauging footprints of positive selection in the genome, or lack thereof, can be informative about the underlying adaptive dynamics^16^.

To elucidate the adaptive dynamics of evolving genomes, it is imperative to determine the genetic composition of evolving populations with high resolution and accuracy^17,18^. Efforts to generate such data from natural populations are often hindered by the inherent challenges of data acquisition and interpretation^19–21^. Long-term evolution experiments (LTEEs) offer a powerful alternative^22^. LTEEs enable us to replicate the evolution of biological (micro-)organisms in simplified, controlled environments^23^. Combined with advancing next-generation sequencing technologies, this framework promises to yield time-resolved genomic data, necessary to study the genetic basis of adaptation at finer resolution^24–26^. Furthermore, by employing different microorganisms, LTEEs allow us to systematically investigate potentially universal links between sequence evolution and adaptation.

Here, we describe in detail the dynamics of the genome evolution of human immunodeficiency virus type-1 (HIV-1) during the first five years of an LTEE. HIV-1 is a retrovirus renowned for rapid adaptation to diverse selective pressures, including host immune responses and antiviral drugs^27,28^. Its efficient adaptability is mainly attributed to its exceptionally high mutation rate (10^-4^-10^-5^ mutations per nucleotide per replication^29^), large population sizes^30^, and relatively short generation time (1.2 days^31^). These features of the virus are also expected to facilitate rapid evolution under experimental settings, such as those in our cell cultures^32^. In addition to high adaptability, HIV-1’s compact ∼9.2 kb genome means that the virus can explore a sizable fraction of its fitness landscape within practical experimental time frames. In conjunction with a well-annotated genome and rich evolutionary literature, HIV-1 is therefore an exceptional model organism for studying the changing dynamics of adaptation based on genome evolution in a stable environment.

This work intends to build upon Bertels *et al.* (2019)^32^, which reported striking levels of parallel evolution in four independently evolving HIV-1 populations using samples from the first 90 transfers of the same experiment. Here, we include additional sequencing and experimental data up to transfer 500, equivalent to ∼1,500 generations and ∼4.8 years of viral passaging, representing the longest-running evolution experiment of HIV-1. The analysis of the extended dataset uncovers a shift from increasing to decreasing parallelism across all populations. As parallel mutations that reach fixation are very likely adaptive^26,32–35^—a premise we support with multiple lines of evidence in our data—this transition marks a swift and substantial deceleration in adaptation. We ascribe this deceleration, at least partly, to the diminishing effects of adaptive mutations. Notably, we also describe genomic patterns consistent with a hard selective sweep in one population at later stages of the experiment, underscoring that episodes of adaptation may occur even after its deceleration. To promote further research, the curated genomic data can be accessed and explored via LTEEviz, an interactive web-based application.

## Results

We passaged the ancestral HIV-1_NL4-3_ strain in the absence of immune responses or antiviral drugs in two different human T-cell lines (MT-2 and MT-4), with two replicates each (Fig. 1). These two cell lines are both HTLV-1-transformed, human leukemic T-cells, but are derived from two different donors and are associated with slightly different characteristics^36^. In each evolution line, 2 µl of the previous infected cell suspension was transferred into a new flask with fresh cell culture twice per week (500 times in total). Every 10th transfer, the genomic composition of populations was determined by next-generation sequencing for each of the four evolutionary lines. Raw sequencing reads were processed via a data pipeline (supplementary fig. S1).

**Fig. 1.**
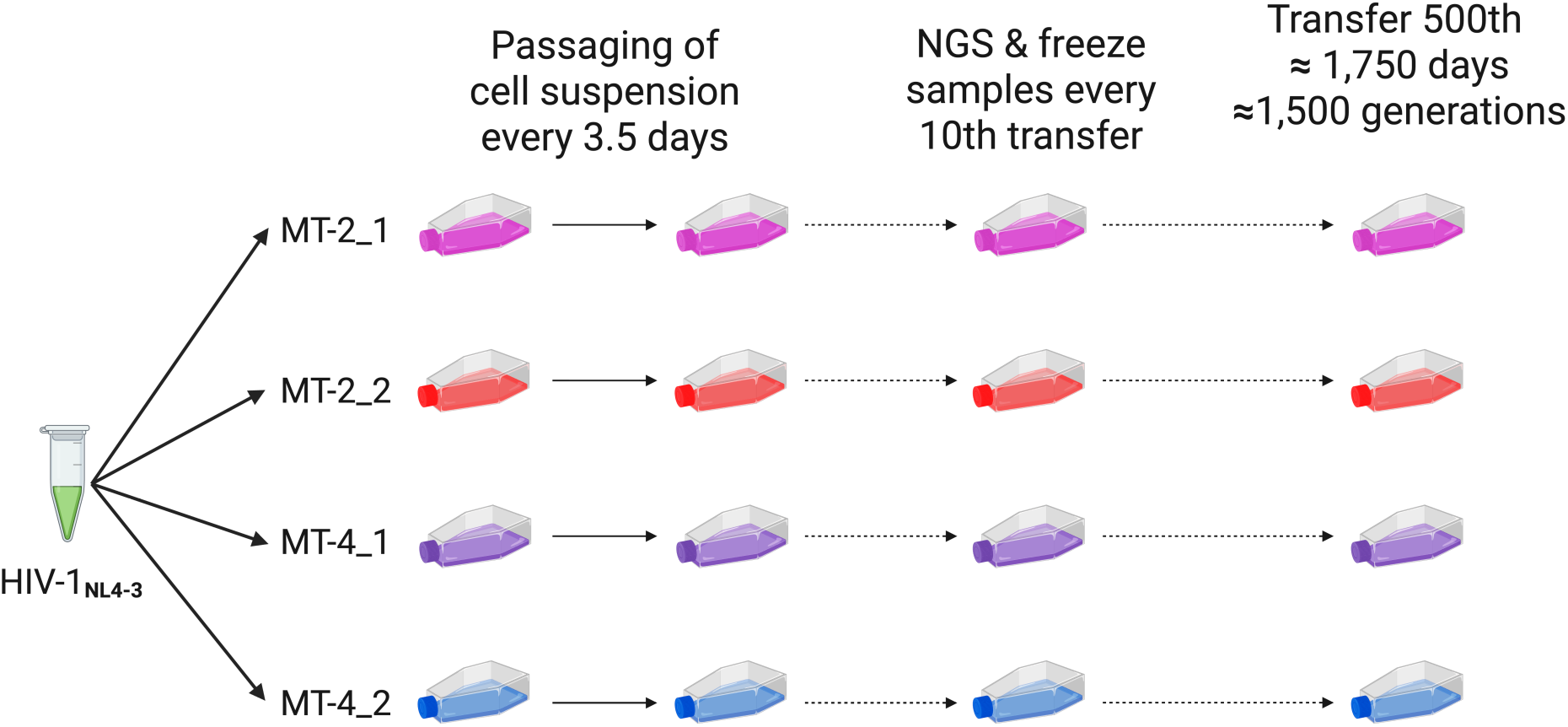
Overview of long-term evolution experiment. The ancestral virus HIV-1_NL4-3_ strain was passaged in four independent lines, with two biological replicates in each of two distinct human leukemia T-cell lines, MT-2 and MT-4. Every 3 or 4 days, 2 µl of infected cell suspension were transferred to anew flask containing uninfected cell culture. At every 10th transfer, cell-free supernatant of infected cell cultures were collected and used for full-length genome sequencing of evolving HIV-1 populations.

We performed titration assays and qPCR to measure the key biological properties of the evolving HIV-1 populations. The multiplicity of infection (MOI) of each transfer for MT-2 lines ranged from 0.00018 to 0.0072 (median of 0.0016) and for MT-4 lines from 0.00039 to 0.027 (median of 0.0021) (supplementary fig. S2). Since each fresh cell culture contained ∼400,000 uninfected T-cells, that roughly translates to a bottleneck size of 639 [IQR: 369,1109] and 842 [IQR: 369,2533] infectious units in 2 µl of inoculum transferred in MT-2 and MT-4 lines, respectively. In concordance with lower overall MOI in MT-2, effective population size estimates (*N_e_*) were significantly lower for MT-2 compared to MT-4 lines throughout the experiment (supplementary fig. S3, mean of 2.8 vs. 3.6 on log10 base).

Additionally, qPCR results show no considerable HIV-1 population growth or death throughout the experiment (supplementary fig. S4, except for MT-2_1). Given a bottleneck fraction of 0.05%, we therefore assumed that HIV-1 populations grew 2,000-fold between each subsequent transfer. As previously justified in Bertels *et al.* (2019), we expect the infective viral offspring number (*R0*) to fall between 12 and 44. Since the measurements of *R0* for HIV-1 both in vivo^37^ and in cell culture^38^ are below the lower bound of this estimated range, we consider an *R0 of 12* for our HIV-1 populations. An *R0* of 12 in turn mandates the completion of three generations during each incubation period of ∼3.5 days, resulting in a generation time of ∼1.2 days, which is within the range reported for HIV-1^39^. Furthermore, the harmonic mean of census population sizes for MT-2 and MT-4 throughout each incubation period would be 4.37 and 5.06 on log10 scale (ratio of 4.8). Therefore, based on these calculations and considering a half-life of 1.6 days for infected cells^40^, the MOI is expected to approach 1 towards the end of the incubation period, thereby increasing the chance of coinfection and hence recombination.

### Increase in Genetic Diversity Is Slowing, Reaching Equilibrium in MT-2 Populations

To investigate the genome evolution of the four HIV-1 populations, we first quantified the genetic diversity using Shannon’s index. The rate of increase in genetic diversity was initially comparable across hosts during the first 60 transfers of the experiment (Fig. 2A). However, the rate began to decline earlier in MT-2 than MT-4 lines. MT-4 populations continued to accumulate genetic diversity even towards the end of experiment, albeit at a reduced rate after transfer 120. In contrast, diversity levels in MT-2 lines nearly plateaued at around H = 0.022 bits by transfer 400.

**Fig. 2.**
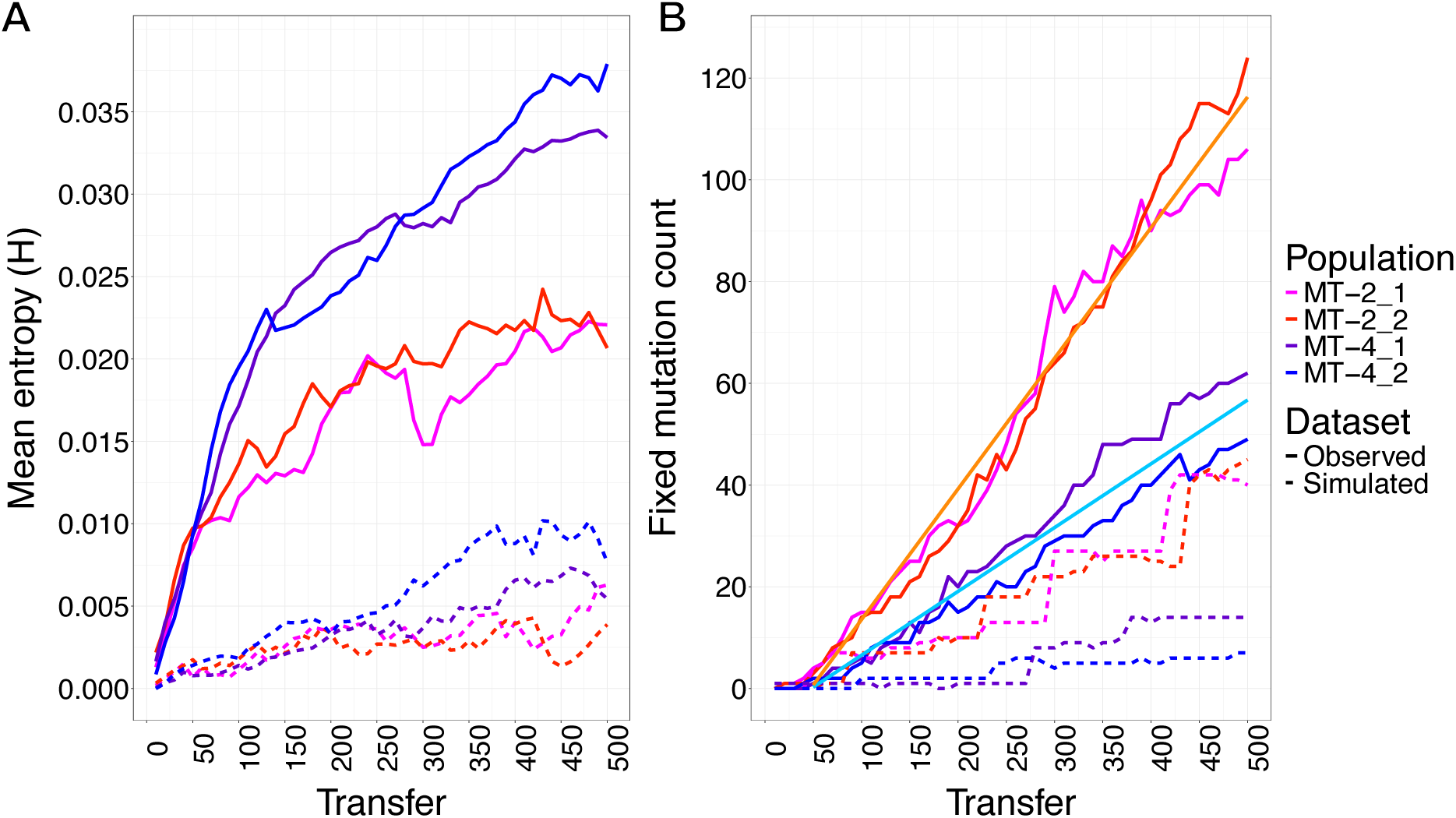
Genomic patterns of diversity and divergence. (**A**) Genetic diversity over time. Population-level genetic diversity was measured per nucleotide by the Shannon’s index and averaged over all the positions. The Shannon entropy per site quantifies uncertainty in the base composition of a randomly chosen genome. For DNA sequences, it ranges from H = 0 meaning the base nucleotide of a site is identical across genomes (only one nucleotide present) to H = 2 bits meaning all 4 nucleotides are equally frequent (*P* = 0.25 each) at a given site. The mean diversity for a sample was then calculated by averaging Shannon indices over the length of the genome. Diversity of all lines increase similarly over the first 60 transfers, but afterwards the rate slows down earlier for MT-2 lines. Moreover, diversity levels for MT-2 lines fluctuate more than MT-4 lines, indicating less stability in the MT-2 populations. Shannon diversity of simulated neutrally evolving populations reflected the observed patterns (dashed lines, refer to Methods). **B**) Count of fixed mutations over time. Mutations continued to fix (frequency ≥ 0.99) at an approximately constant rate even towards the end of experiment. There were no significant deviations from linearity (ANOVA, *P* > 0.05). The four evolution lines harbored 106, 124, 62, and 49 fixed mutations at transfer 500, respectively. The extrapolated slopes of 0.256 and 0.125 (fixed mutations per transfer) for MT-2 and MT-4 lines are shown in orange and light blue, respectively. Simulation of neutral sequence evolution (dashed lines, refer to Methods) suggests that 37% (∼43 mutations) in MT-2 and 19% (∼11 mutations) in MT-4 of the observed fixed mutations are expected to arise by neutral evolution alone.

Genetic diversity reaches an equilibrium when mutation and drift are in a balance^41^. In agreement with smaller effective population size estimates for MT-2 lines, we expect that stronger drift to be responsible for lower standing genetic variation observed in MT-2. First, similar rates of diversity increase during the first 60 transfers suggest that the mutation rates in HIV-1 populations, at least initially, were comparable between the two hosts. This is because early in the experiment, when there exists lower overall diversity, genetic diversity can be approximated to be proportional only to mutation input^42^. Second, we observed a significantly higher rate of diversity loss in MT-2 lines compared to MT-4 (*P* < 0.001, paired t-test; supplementary fig. S5), reflected by a greater number of minority mutations (0.01 ≤ frequency < 0.5) disappearing between each consecutive samples.

### Mutations Continue to Fix at Approximately Constant but Host-Specific Rates

Next, we analyzed the temporal patterns of fixed mutations. We define mutations as “fixed” if they are present at a frequency of 99% or more in the population of a given sample. In total, we identified 414 single-base substitutions and 2 short deletions that reached fixation in either of four lines at some point during the experiment. The strong bias towards single-base substitutions over indels in the HIV-1 genome evolution is well-documented in both patient and cell culture studies, and is attributed to a combination of mutational mechanisms and selective pressures^43–45^.

Mutations were fixed at an approximately constant rate throughout the experiment in each population (Fig. 2B). The ongoing near-linear fixation of mutations despite saturation of genetic diversity resembles patterns of evolving HIV-1 populations in treatment-naïve patients^46,47^. The rate at which mutations fix can be calculated as the slopes in Fig. 2B. Assuming a generation time of 1.2 days, the fixation rates are 0.085 and 0.042 per generation for MT-2 and MT-4 lines, respectively. Therefore, mutations fixed virtually twice as fast in MT-2 (*P* < 0.001).

To pinpoint the source of the extra fixed mutations in MT-2 lines, we categorized them based on their translational impact. Fixed mutations were most densely distributed in non-coding regions (supplementary fig. S6); but, the ratio of coding to non-coding fixed mutations for MT-2 and MT-4 lines was not significantly different from each other. In protein coding regions, however, we observed significantly larger ratio of synonymous to nonsynonymous fixed mutation in MT-2 lines. While the number of synonymous fixed mutations per synonymous site was significantly higher than that of nonsynonymous in every line (*P* < 0.001, paired t-test; supplementary fig. S7), the effect size of this difference was considerably larger in MT-2 compared to MT-4 lines (Cohen’s *d* = 1.18 vs. 0.54, respectively). These results suggest that the excess of fixed mutations in MT-2 lines predominantly came from synonymous mutations.

### High Parallelism Observed Among Fixed Mutations

Parallel evolution is the independent evolution of identical genetic changes and can arise when the same adaptive solutions are repeatedly favored by selection^48^. Therefore, parallelism at the nucleotide level is often used to identify potential targets of selection and gauge its strength^26,33^. We measured parallelism among fixed mutations across evolutionary lines. As all our evolutionary lines were initiated from a genetically homogenous ancestral population, the significance of observing parallel fixed mutations is that these genetic changes had to first independently arise by mutation in each evolution line and then be driven to fixation by selection.

Fig. 3A shows a Venn diagram of fixed mutations present at transfer 500 and highlights considerable parallelism among different lines. Overall, 51 out of 274 distinct fixed mutations at transfer 500 (18.6%) evolved and fixed independently in at least two evolution lines, and 13 mutations (4.7%) in at least three lines. Notably, three fixed mutations emerged and fixed among all four lines. Two of these occurred in the 5′ untranslated region (5′UTR) and 5’ long terminal repeats leading sequence (5′LTRLS) of the HIV-1 genome (G566A and G678A). The third mutation was a nonsynonymous mutation in *env* (A7854G or Asp545Gly). This Asp-to-Gly substitution is anticipated to have a low biochemical distance (Grantham score of 56^49^), suggesting it is unlikely to disrupt *env* structure or function. This mutation represents a reversion to the so-called HIV-1 “consensus of consensus” sequence compiled from all HIV-1 genomes in the Los Alamos database.

**Fig. 3.**
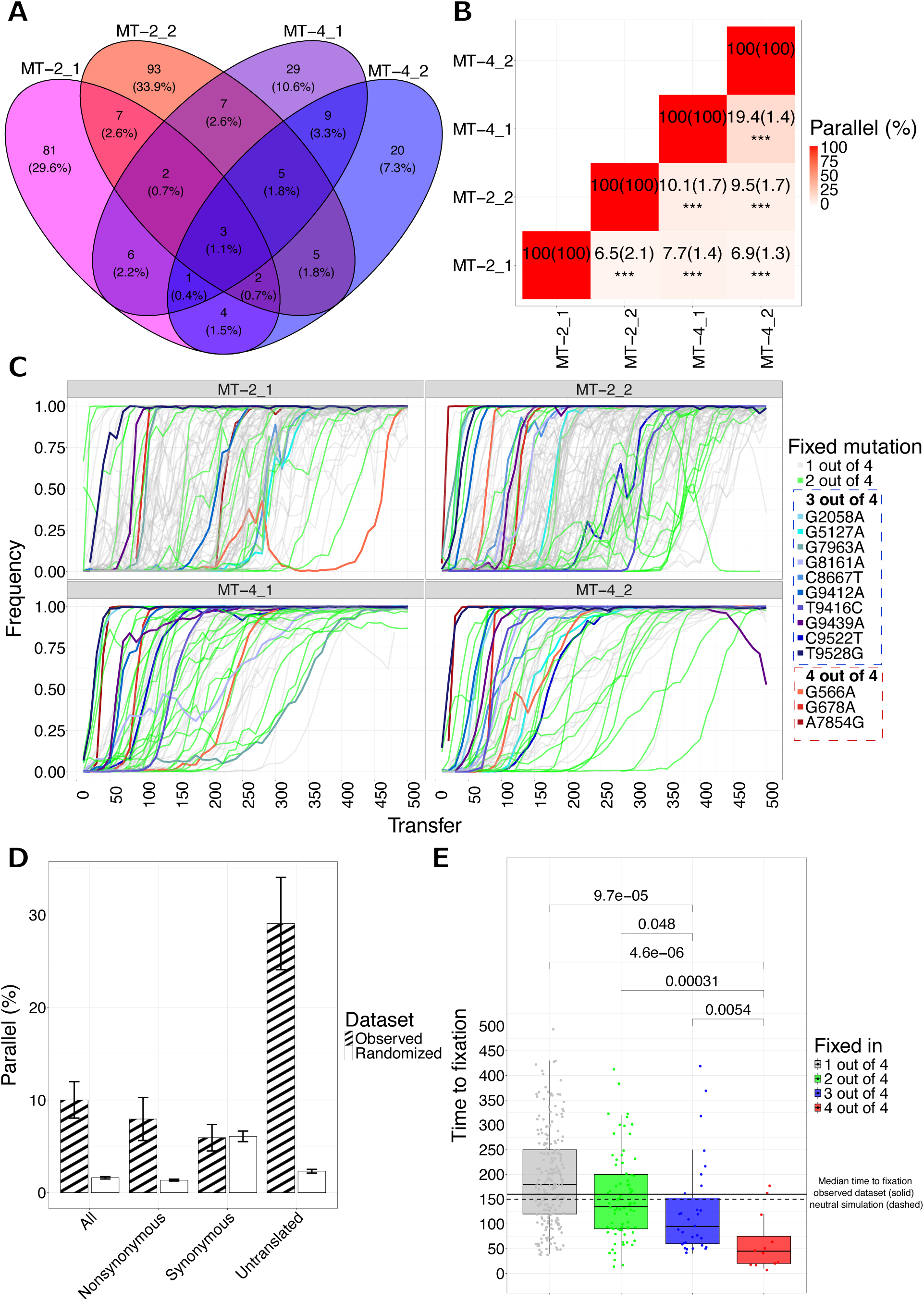
Abundance and characteristics of parallel fixed mutations. (**A**) Venn diagram depicting number of shared and unique fixed mutations at transfer 500. An annotated list of these fixed mutations can be viewed via the data table in LTEEviz. Overall, out of 274 distinct fixed mutations present at passage 500, 223 (81.4%) are found in only a single evolution line. (**B**) Summary heatmap showing percentage of parallel fixed mutations between each two replicates. Numbers in parentheses contain expected level of parallelism computed by permutation tests. In all comparisons, observed parallelism was significantly higher than expected. (**C**) Frequency trajectories of fixed mutations. Mutations fixed in one and two populations are shown by grey and green, respectively. Mutations fixed in three and four populations are shown individually by shades of blue and red, respectively. (**D**) Percentage of parallel fixed mutation among each two lines in observed and randomized datasets per translational impact group. Synonymous fixed mutations were not significantly different than expected, while fixed mutations in nonsynonymous and non-coding regions were significantly more parallel in observed dataset compared to randomized (Cohen’s *d* = 1.6 vs. 3.1, respectively). Error bars represent the mean ± standard error. (**E**) Time-to-fixation of fixed mutations based on their number of lines they were fixed in. Time-to-fixation was defined as the number of transfers it took a mutation to appear above 0.01 frequency and (continuously) reach fixation by surpassing 0.99 frequency. The solid and dashed lines depict the median time-to-fixation from the observed dataset (160) and neutral sequence simulation (150), respectively.

Overall, we observed high levels of genetic parallelism across all lines, but to varying degrees. Fig. 3B summarizes parallelism among fixed mutations between each line. At transfer 500, 19.4% of fixed mutations in MT-4 lines were shared between its two replicates, while this number was 6.5% for MT-2 replicates. Of note, the average level of parallelism between replicates of different environments was 8.55%.

### Observed Parallelism Exceeds the Level Expected Under Neutral Evolution

We carried out permutation tests to determine to what extent the observed levels of parallelism of fixed mutations deviate from the expectation under neutral evolution. The permutation tests entailed generating expected fixations by randomly drawing from the pool of all mutations (with the same reference and alternative alleles) observed above 0.05 frequency in each evolution line at some point throughout the experiment (refer to Methods). By this approach, we accounted for the potential contributions of mutation bias and purifying selection to parallelism. The results of permutation tests (Fig. 3B, numbers in brackets) confirmed that in all lines the observed levels of parallelism were significantly greater than expected under neutral evolution.

Additionally, randomization tests accounting for the cell line indicated that parallelism observed between MT-4 replicates was significantly larger than expected if the fixed mutations were to be randomly redistributed between evolution lines, while the same effect was not seen for MT-2 replicates (supplementary fig. S8). A contributing factor to the reduced parallelism observed in MT-2 relative to MT-4 lines is genetic hitchhiking^50^. In line with signatures of genetic hitchhiking, the frequency trajectories of MT-2 fixed mutations in Fig. 3C appeared to be more synchronized than MT-4 trajectories. More prevalent hitchhiking in MT-2 lines can be the result of stronger linkage. Indeed, direct comparison of linkage disequilibrium confirmed this pattern: MT-2 lines exhibited significantly stronger linkage than MT-4 lines (supplementary fig. S9; paired *t*-test, *P* < 0.001).

We observed two conspicuous examples of genetic hitchhiking in our experiment. In MT-2_1, between transfers 270 and 310, a set of 22 mutations doubled in frequency and reached fixation by transfer 500. Of these mutations, 7 were synonymous: 5 unique to a single line and 2 fixed in two lines. In a second case in MT-2_2, between transfers 160 and 200, a set of 25 mutations likewise doubled in frequency and reached fixation by transfer 500. This set included 12 synonymous mutations: 11 unique to a single line and 1 fixed in two lines. These cases highlight how multiple, probably neutral synonymous mutations can rise in frequency synchronously and fix within individual MT-2 lines yet show little convergence across lines. These observations provide an explanation for our finding above that synonymous mutations account for most of the fixations in MT-2 lines.

### Evidence for Higher Fitness of Parallel Fixed Mutations

In addition to the evidence for excess parallelism from permutation tests, specific characteristics of these mutations further support the key role of selection in driving them to fixation. Firstly, synonymous mutations are underrepresented among parallel fixed mutations. Only 6% of all parallel fixed mutations were synonymous, while around 21% of all possible mutations in HIV-1 genome would lead to a synonymous change. In fact, the level of parallelism in synonymous mutations was not significantly different than what we expect under neutral evolution (Fig. 3D).

Secondly, the frequency trajectories of parallel fixed mutations appeared to rise more sharply depending on how many times they emerged as fixed across different lines (Fig. 3C). To quantify their fixation kinetics, we measured the duration from their appearance to their fixation (time-to-fixation). Parallel fixed mutations exhibited significantly shorter time-to-fixation (*P* < 0.05, Wilcoxon rank-sum test) with a clear trend: the time to fixation decreased with the number of evolution lines in which the mutation occurred (Fig. 3E). For instance, the median time-to-fixation for mutations fixed in all four lines was approximately 45 transfers, compared to 200 transfers for line-specific mutations (those fixed only in one line). The monotonic decrease in time-to-fixation with increasing parallelism strongly supports the role of selection in driving parallel mutations to fixation. As the time to fixation of highly-parallel fixed mutations (those fixed in three or four lines with total of 13 distinct mutations) is significantly lower than median time to fixation of neutral sequence simulation (*P* < 0.01, Wilcoxon rank-sum test), we consider them as “putatively adaptive” and focus on their dynamics in the next section.

### Diminishing-Returns, Not Sign Epistasis, Define Interactions Between Adaptive Mutations

To obtain a complete picture of the evolutionary dynamics in our long-term experiment, we investigated epistatic interaction among adaptive mutations. First, we focus on the most extreme form of epistasis: sign epistasis which reverses the fitness effect of a mutation from positive to negative, or vice versa^51,52^. To test whether sign epistasis had been involved in constraining emergence of adaptive mutations, we inspected the chronological order of their fixation (Fig. 4A). We found that the three mutations that fixed all four lines did so in three different orders across the lines. Additionally, five out of 10 mutations that were fixed in same three lines did not follow any pattern in their order of fixation. A distinct order of fixation per line among adaptive mutations suggests that these mutations were beneficial in different genetic backgrounds.

**Fig. 4.**
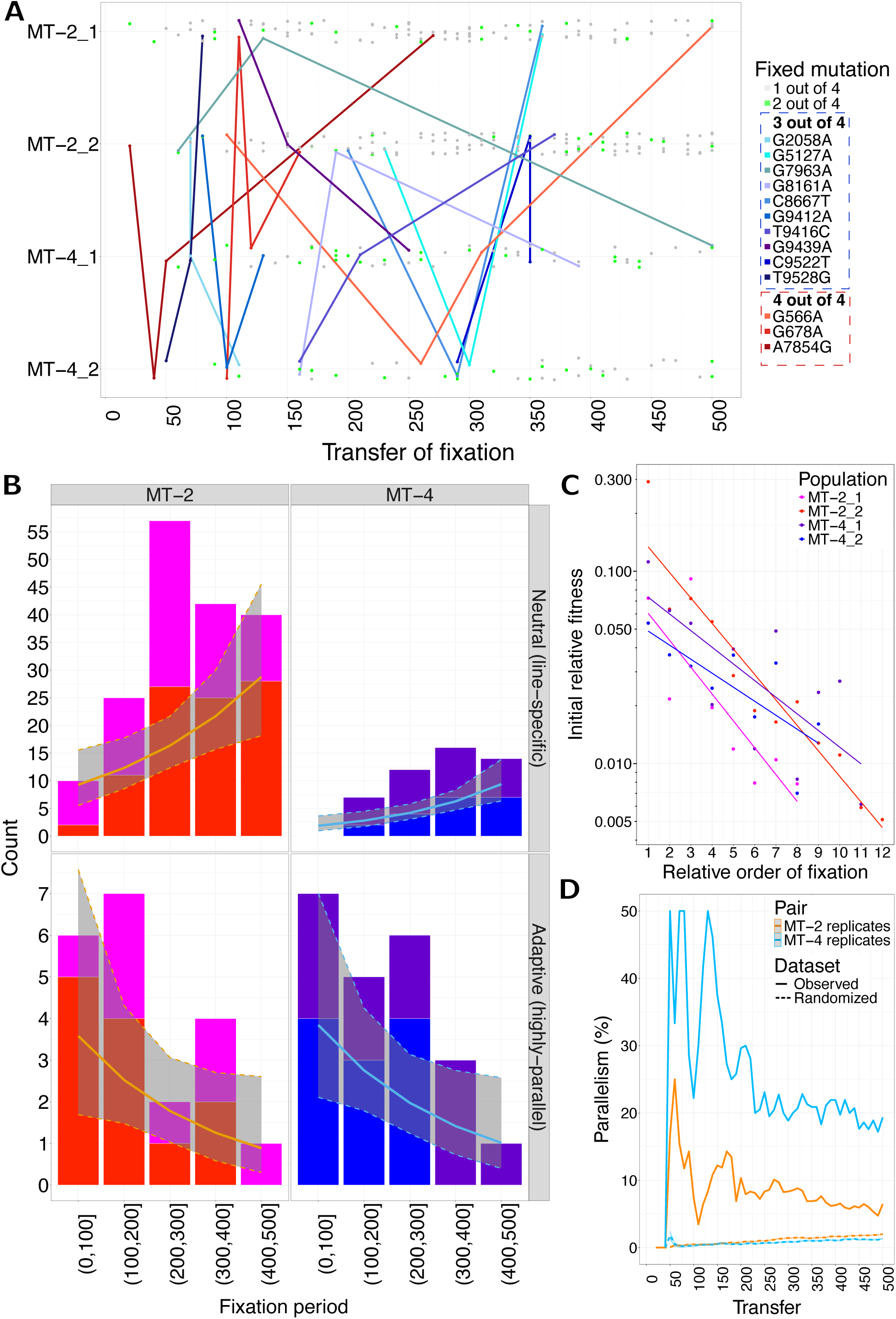
Interactions between parallel mutations and their temporal dynamics. **(A) Order of fixed** mutations based on the transfer at which they fixed. Mutations fixed in one and two populations are depicted by grey and green points, respectively (lines for mutations fixed in two populations are omitted for clarity). Mutations fixed in three and four populations are depicted case by case with points and connecting lines in shades of blue and red, respectively. Out of 10 mutations that fixed in three lines, five (G2058A, G8161A, G9412A, T9416C, C9522T) fixed in the same three lines (MT-2_2, MT-4_1, MT-4_2), but their order of fixation did not contain any pattern. (**B**) Counts of neutral (line-specific) and adaptive (highly-parallel) fixed mutations per their fixation period aggregated for replicates of MT-2 and MT-4. Each bin represents 100 transfers. Notably, the number of neutral (line-specific) mutations is markedly larger in MT-2 lines. (**C**) Initial relative fitness vs. relative order of fixation (time). Each point represents an adaptive mutation colored by the respective line they occurred in. The x-axis represents the order at which each adaptive mutation was fixed relative to other adaptive mutations in the same line. Therefore, x-axis represents relative time specific to each line. All fitted regression lines are significant after multiple testing correction. The magnitude of decline tended to be greater in MT-2 than in MT-4 lines (average slope estimate of -0.471 vs. -0.251), but the difference was not statistically significant between the two host environments. (**D**) Parallelism among fixed mutations between replicates of MT-2 and MT-4. The neutral sequence simulation (dashed lines) estimates increasing expected parallelism among fixed mutations over time (ribbons indicate lower and upper 95% CI). Conversely, the observed level of parallelism is generally decreasing since transfer 60 and 130 in MT-2 and MT-4 replicates, respectively. The decreasing parallelism indicates positive selection during the beginning of the experiment was stronger.

We next tested whether the magnitude of fitness effects of adaptive mutations was modulated by the genetic background, without changing their sign. To quantify the mutational effects, we used the “initial relative fitness” metric proposed by Lang *et al.* (2011)^53^. The premise of this measurement is that the initial frequency increase of a mutation should be proportional to the selective advantage it provides in its genetic context relative to the population average. Consistent with previous findings,^25^ logistic regression modeling confirmed that probability of fixation of mutations was partly explained by their initial relative fitness (supplementary fig. S10).

To test the temporal trends of fitness effects of adaptive mutations, we fitted linear regression models for initial relative fitness of adaptive mutation as a function of time for each evolution line separately (Fig. 4B). All four models confirmed a significant (adj.*P* < 0.05) negative log-linear relationship between initial relative fitness and time (represented as the relative order of fixation of adaptive mutations relative to each other within a single line).

Based on models’ slope estimates, each adaptive mutation was associated with 27.8% (MT-2) and 15.9% (MT-4) lower initial relative fitness compared to the previous adaptive mutation in the same line. In other words, 90% of projected fitness gain by these adaptive mutations was achieved within 260 and 310 transfers (∼720 and ∼930 generations) in MT-2 and MT-4 populations, respectively (supplementary fig. S11). Additionally, we found that the initial relative fitness of the same adaptive mutation was 1.89 times larger (median) in the evolution line in which it went to fixation earlier (*P* < 0.05, Wilcoxon rank-sum test, supplementary fig. S12). These results highlight the presence of strong and pervasive diminishing-returns epistasis in our HIV-1 populations, causing adaptive mutations to confer consistently smaller fitness gains in fitter backgrounds.

Moreover, accumulation of adaptive mutations slowed down considerably over the course of the experiment (Fig. 4C). We utilized generalized linear mixed-effects models to quantify the downtrend in counts of adaptive mutations and contrast it to that of neutral mutations. Our models confirmed that the count of adaptive (highly-parallel) mutations significantly decreased over time by multiplicative factors of 0.57 and 0.59 per 100 transfers in MT-2 and MT-4 populations, respectively. This pattern likely results from a limited supply of beneficial mutations with large effects and their progressively shrinking beneficial effects due to the influence of diminishing-returns epistasis. In contrast, line-specific fixed mutations increased significantly over time, rising by multiplicative factors of 1.56 and 1.9 per 100 transfers in MT-2 and MT-4, respectively. Because these line-specific mutations are likely neutral and fixed through genetic drift and hitchhiking, this pattern reflects the ongoing activity of stochastic processes. Overall, our models strongly support an opposite temporal trend in fixation rate of adaptive (highly-parallel) and neutral (line-specific) fixed mutations in both hosts (*P* < 0.01, refer to supplementary fig. S13 for model results and performance).

### Parallelism Decreases as Populations Continue to Evolve

Bertels *et al.* (2019) also noted diminishing fitness effects among majority mutations (frequency ≥ 0.5) by passage 90 but did not report a corresponding reduction in parallelism. We hypothesized that, over longer timescales, a decrease in parallelism might be observable. This expectation is grounded in not only the declining supply of adaptive mutations with progressively smaller fitness gains as we showed, but also in the constant activity of drift fixing neutral mutations.

To test this hypothesis, we quantified the proportion of parallel fixed mutations relative to all fixed mutations between replicates of each host over time (Fig. 4D). The proportion of parallel fixed mutations among MT-2 evolution lines peaked at 25% at transfer 60 (2 out of 8 mutations) and dropped to 6.5% by transfer 500 (14 out of 216 mutations). Similarly for MT-4 evolution lines, parallelism peaked at 50% at transfer 130 (6 out of 12 mutations) and dropped to 19.4% by transfer 500 (18 out of 93 mutations). The time trends of proportion of parallel fixed mutations was congruent to that of the rate of nonsynonymous to synonymous fixed mutation per site (supplementary fig. S14). By contrast, permutation tests indicated that, under neutral evolution, parallelism is expected to gradually increase over time (Fig. 4D). This pattern emerges as random processes continue to fix mutations across a finite set of (nearly) neutral genomic loci.

### A Selective Sweep Occurs After Deceleration of Adaptation in MT-2_1

Patterns of genomic diversity and divergence support the occurrence of a (hard) selective sweep in MT-2_1 from transfer 270 to 310, after the adaptation had decelerated substantially in this evolution line. First, during this period, the rate of divergence in MT-2_1 was increased significantly as manifested in the reconstructed phylogeny (supplementary fig. S15). Second, this period coincided with a 20% drop in diversity, followed by a rebound to equilibrium levels (Fig. 2A). The diversity reduction is one of the main genetic hallmarks of (hard) sweeps^54,55^, which is caused by a single genotype outcompeting every other genotype and purging polymorphism surrounding the site(s) undergoing selection.

Identifying the driver of this selective sweep with the available data is challenging, but several cases among the set of 22 mutations that doubled in frequency during this period and reached fixation by transfer 500 are worth noting. The first case is the emergence of a rare amino acid change in *gag*. Two mutations whose frequencies rose sharply and in a synchronized manner during this period occurred at adjacent positions 973, and 974 within the *gag* gene (Fig. 5A). These two mutations are nonsynonymous and affect the same codon (codon 62 of the 500-residue *gag* MA protein), with each mutation leading to a different amino acid change if it occurs independently. Frequency trajectories of these mutations indicate that G974C appeared first, resulting in Gly62Ala amino acid change, followed by the emergence G973A. Based on inspection of raw sequencing reads, G973A occurred in the background of the G974C, leading to a subsequent Ala62Thr amino acid change at the same position of the amino acid. Of note, there was no single nucleotide change in that codon that could have resulted in direct Gly → Thr amino acid change at position 62 of *gag* MA protein. Moreover, both G973A and G974C were not observed above frequency of 0.05 in any other line than MT-2_1. This observation suggests that each of these mutations separately could have been deleterious for the virus, while their co-emergence reversed their mutational effect (reciprocal sign epistasis) from deleterious to beneficial (or neutral), and thereby facilitated their fixation in MT-2_1.

**Fig. 5.**
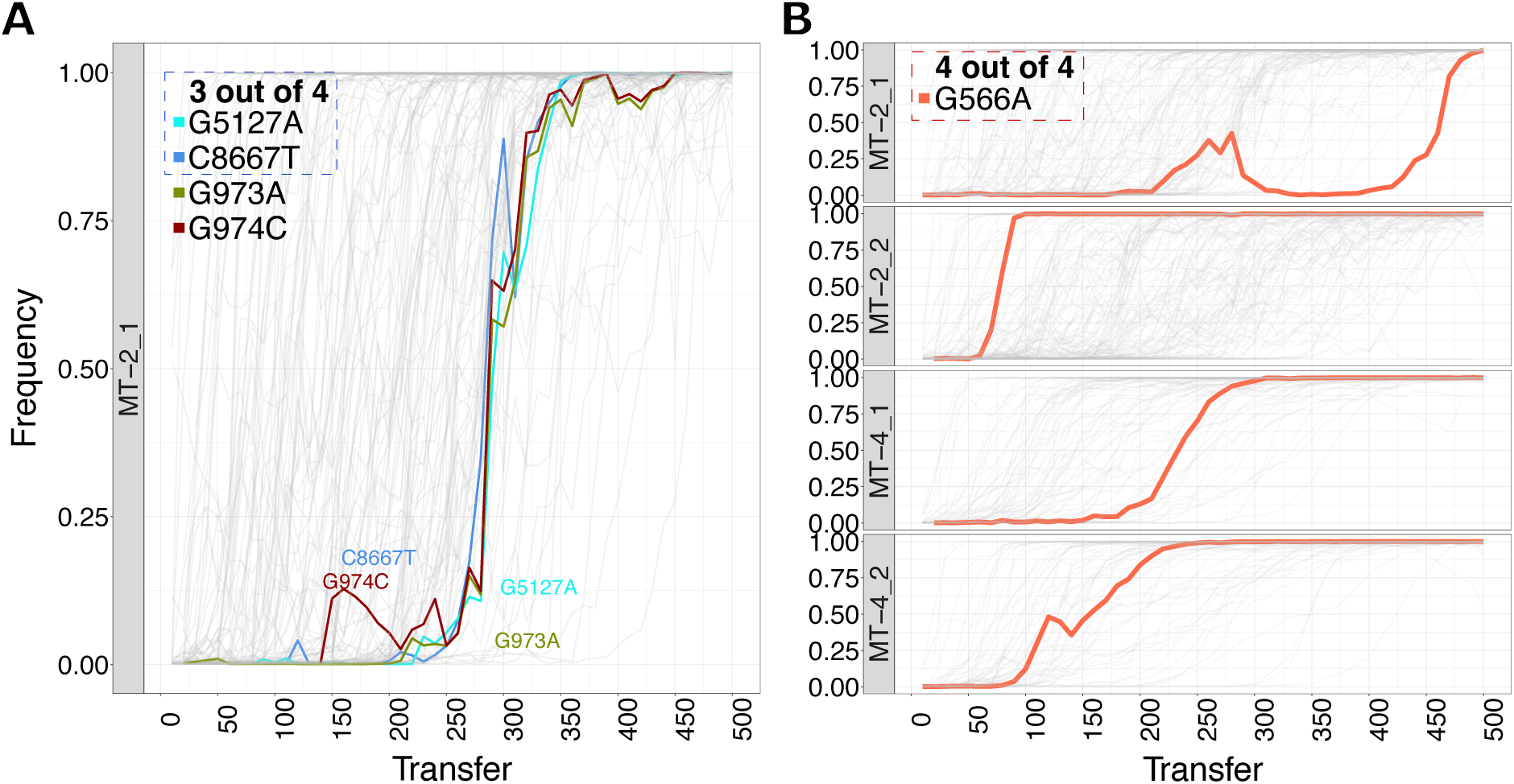
Selective sweep and clonal interference in MT-2_1. (**A**) Frequency trajectories of potential drivers of the selective sweep. Although we lack haplotype information linking these sites together, their tightly correlated trajectories, together with a 20% drop in global diversity despite ongoing recombination, suggest that they could have resided within the same sweeping genotype. (**B**) An example of clonal interference between the sweeping genotype and another genotype with a putatively beneficial mutation in MT-2_1. G566A emerged and rose to fixation continuously and through a straightforward trajectory in all evolution lines, but in MT-2_1. In MT-2_1, its frequency trajectory was interrupted by the selective sweep. After being eliminated from the population following the sweep, G566A re-emerged at transfer 410 and reached fixation by transfer 500 in MT-2_1 population.

Additionally, two adaptive mutations (G5127A and C8667T) also rose synchronously with the sweep. Both mutations were nonsynonymous: G5127A hit *vif*, causing a Met29Ile amino acid change; while C8667T hit *env*, resulting in Thr816Ile. Whether any of these mutations were the driver of this sweep or not cannot be inferred from our data, but their presence in the outcompeting genotype could provide further fitness advantage over the existing genotypes in the population.

Independent evidence in support of the fitness benefit of this sweeping genotype comes from its clonal interference with another genotype in MT-2_1, which carries the beneficial mutation G566A (Fig. 5B). G566A was fixed in all four lines with average time-to-fixation of 125 transfers compared to the overall average of 170 transfers, pointing to its potential benefit for the fitness of the virus. Its frequency paths to fixation were straightforward in all lines, except in MT-2_1. In MT-2_1, G566A’s frequency trajectory was interrupted by the occurrence of the selective sweep: it appeared at transfer 180 and rose to around 0.44 frequency by transfer 280, when it was overtaken by the sweeping genotype and disappeared from the population. Later in the experiment, it reappeared by an independent de-novo mutation event and reached fixation by transfer 500. This observation emphasizes the adaptiveness of the sweep and the competition that exist between sub-populations^25^.

## Discussion

In this study, we investigated the adaptive dynamics of HIV-1 based on its genome evolution in a long-term evolution experiment. We observed excess parallel evolution of fixed mutations and provided three independent lines of evidence to establish their adaptive nature. First, these mutations were highly unlikely to be parallel under neutral evolution based on randomization test results. Second, they were significantly underrepresented among synonymous mutations. Third, their time to fixation was significantly lower than that of line-specific fixed mutations or fixed mutations in the neutral evolution simulation.

In the absence of intentionally imposed selective pressures in our study (e.g., immune response or antiretroviral drugs), we expect acclimatization to distinct cell line hosts and in vitro growth conditions to represent the primary source of selective pressure. By examining the prevalence and epistatic interactions among the adaptive mutations, we showed that the rate of adaptation decelerated substantially over the course of the experiment: the fixation rate of adaptive mutations almost halved within the first 300 transfers, while their conferred fitness gain decreased by 27% (MT-2) and 18% (MT-4) per every added adaptive mutation in their genetic background. This deceleration was also reflected in infectivity data obtained via titration assays: in all evolutionary lines, we detected an initial phase of increasing infectivity up to transfer 150 in MT-2 and transfer 300 in MT-4 populations, after which infectivity levels stabilized.

Previous studies documented the deceleration of adaptation of evolving microbial populations in constant environments for a diverse set of microorganisms such as bacteria^26^, phage^56^, and yeast^3^, indicating that this phenomenon transcends the specifics of the biological system^14^. Here, we highlight that the timing and scale of such deceleration can vary greatly based on the molecular and demographic properties of the model organism^57,58^. In *E. coli*, for instance, Tenaillon *et al.* (2016)^26^ reported that “the proportion of observed mutations that were beneficial declined over time but remained substantial even after 50,000 generations”. By contrast, our analysis of HIV-1 genome evolution indicates that a substantial decline in adaptation was reached within the course of our experiment, which corresponds to 1,500 HIV-1 generations. To quantitatively compare the two experiments, we relied on the measurement of the rate of nonsynonymous to synonymous mutations reported in Tenaillon *et al.* (2016). In *E. coli* populations, this rate dropped to two at generation 50,000, while in our HIV-1 populations the rate of two was reached by transfer 70 (generation ∼210) and 230 (generation ∼690) in MT-2 and MT-4, respectively.

Several factors could underlie earlier deceleration of adaptation in our HIV-1 populations compared to that of *E. coli* populations. Most important among those is HIV-1’s enhanced ability to diversify due to higher mutation rate and possibility of recombination compared to *E. coli*. Mutations are the fuel for selection; the faster adaptive mutations appear, the earlier they will get fixed in the population by natural selection. However, despite the mutation rate estimates of HIV-1 being several orders of magnitude higher (∼2 × 10^-5^ mutations per nucleotide per replication^29^ vs. 8.9 × 10^−11^ per base-pair per generation in non-mutator lineages of *E. coli*^59^), the population mutation supply (*μ* × *N_e_*) was only around two orders of magnitude larger for HIV-1 populations (3.17 × 10^-2^ vs. 8.9 × 10^-4^). To what extent does this difference in mutational supply explain the timescale of the deceleration of adaptation will need to be answered by detailed population genetic simulation studies in the future.

Recombination can also increase the rate of appearance of new genotypes through random shuffling of the genetic material between different haplotypes^60^. In doing so, it alleviates the clonal competition between sub-populations by increasing the efficacy of selection (the Fisher-Muller effect^61–63^). While the reproduction of *E. coli* strain was strictly clonal (i.e., no horizontal gene transfer^64^), HIV-1 can recombine through template switching^65^, resulting in faster adaptation over a wide range of conditions as shown previously^66^. The occurrence of recombination in our evolving HIV-1 populations was evident from the fast decay in linkage disequilibrium of mutations within the span of read pairs.

Notwithstanding the substantial decline in rate of adaptation, we observed that all populations continued to diverge from the ancestor at a relatively constant rate even towards the end of the experiment. Such prolonged period of linear accumulation of fixed mutations has been observed in microbial populations in similar studies^25,67^. This continuing accumulation can be, in part, attributed to the steady fixation of (nearly) neutral mutations via genetic drift^26^. Our simulations of neutral sequence evolution estimated that 37% (∼43 mutations) in MT-2 and 19% (∼11 mutations) in MT-4 lines could have been fixed by drift alone. The sources of drift in our experiment are periodic bottlenecking, variation in *R0* among individual infections, and founder effects due to finite population sizes. But resolving the discrepancy between the linear accumulation of fixations on the one hand, and the deceleration of adaptation, on the other, will require further investigation.

Two limitations of our study are worth noting. First, although we substantiated the role of selection in giving rise to parallelism among fixed mutations across evolutionary lines, we did not disentangle the potential contributions of biased mutation rates to the extent of observed parallel evolution^68–71^. Nonetheless, we believe the undesired influence of mutational bias should be mitigated by focusing on parallelism at the level of fixed mutations. The second limitation, inherent to the use of short-read pooled sequencing data, is the lack of haplotype information. Available computational tools failed in reconstructing haplotypes, restricting us from thoroughly examining clonal dynamics within populations and interactions between mutations. As a workaround, we resorted to identifying and characterizing instances of such phenomena using frequency trajectories and the partial linkage information that was available within the short sequencing read pairs. That said, the availability of frozen samples offers a promising avenue for future studies to obtain single-read coverage of the entire HIV-1 genome using long-read sequencing technologies^72,73^.

High-throughput sequencing datasets such as those generated in this study demand extensive preprocessing such as quality filtering, read mapping, and downstream data handling before they can effectively inform specific biological questions. As a result, although public deposition of raw sequencing data is essential, these files alone often have limited practical value for secondary analyses. To enhance accessibility, ensure transparency, and promote broader reuse, we created an interactive web-based platform (LTEEviz) that enables users to examine and visualize the dataset dynamically. The platform also offers straightforward access to the curated dataset, lowering the technical barrier for reanalysis. We anticipate that this resource will support further investigation and enable new insights beyond the scope of the present study.

In summary, our results revealed a swift and substantial decline in the rate of adaptation in our HIV-1 populations. We propose fast diversification as a key driver of such rapid adaptation in HIV-1 compared to that of other microbial populations in similar studies. Future controlled laboratory experiments or computer simulations are needed to model how diversification rates, along with other potentially relevant biological determinants, influence the timing and extent of deceleration in adaptation of microbial populations.

## Material and Methods

### Passaging of HIV-1

The detailed description of HIV-1 passaging in this LTEE study can be found in our previous publication^32^. In summary (refer to Fig. 1A for a graphical overview), the ancestral virus HIV-1_NL4-3_ strain (generated and characterized as previously indicated^74^) was passaged in four independent evolutionary lines in two biological replicates of human leukemia T-cell cell lines^75^, MT-2 and MT-4. The HIV-1 full-length plasmid pNL4-3 and the two cell lines were obtained through the AIDS Research and Reference Reagent Program, Division of AIDS, NIAID, NIH from Dr. Malcolm Martin^76^ and Dr. Douglas Richman, respectively.

At every 3rd or 4th day, 2 µl of infected cell suspension was transferred to a new flask containing 4×10^5^ uninfected cells. At every 10th transfer, cell-free supernatant of infected cell suspension was obtained through centrifugation and stored at -80°C freezer. Of note, every few months a fresh batch of cells were thawed and used for the continuation of the experiment to avoid potential confounder effects associated with co-evolution of cell lines with viral lines.

### Sequencing of HIV-1 Genome

Full-length genome of the virus stock HIV-1_NL4-3_ (ancestor) and every 10th transfer until transfer of 500 (4 × 50 = 200, in total) were sequenced using the Illumina MiSeq next-generation sequencing platform (Illumina, San Diego, California, United States of America) as previously described^74^. This Briefly, HIV-1 RNA was isolated from 150 µl virus stock HIV-1_NL4-3_ or cell-free supernatant. Five overlapping amplicons were generated by reverse transcription polymerase chain reaction covering the full genome of HIV-1 per sample. The 5 amplicons per sample were pooled, and libraries were prepared with the Nextera^®^ XT DNA Sample Preparation Kit (Illumina) according to the manufacturer’s description. Next-generation sequencing was performed using Illumina’s MiSeq Sequencer, generating paired-end reads of 2 × 250 bp length (v2 kit). To minimize the risk of cross contamination, samples from each replicate line were processed separately.

### NGS Data Processing

All the steps of processing raw sequencing reads were implemented and performed via a Snakemake pipeline (supplementary fig. S16 data processing workflow; supplementary fig. S1 pipeline’s directed acyclic graph). First, using FASTQC and MULTIQC tools, we examined the quality of raw sequencing reads. Sequencing samples that did not meet quality requirements were re-sequenced. Next, using Trimmomatic we trimmed base calls with low quality (Phred score ≤20, corresponding to 0.01 error probability) and removed short reads (length of ≤ 36 bases).

The trimmed reads were then mapped against the ancestral HIV-1_NL4-3_ consensus sequence using BWA-MEM with default settings. Reads with mapping quality ≤ 35 (corresponding to error probability of 0.00032) were discarded. The average sequencing depth each of the 5 amplicons were calculated and if they fell below 1,000, the sequencing was repeated for that amplicon. The R-to-R viral genome of HIV-1 for all samples had at least a coverage of 200 quality reads, resulting in full-length sequencing of genome (for coverage summaries refer to supplementary fig. S17).

To call the variants, we used in-house python scripts (99% similarity with commonly used tools such as smalt-align and mpileup). The genome coordinates are based on NL4-3 plasmid (NCBI accession number: AF324493.2). A minor allele frequency (MAF) cutoff of 0.01 and sequencing depth cutoff of 200 reads were applied to exclude potential erroneous sequencing calls. For downstream analyses, mutation categories were defined based on the following variant frequency ranges:

| Mutation Category | Frequency |
| --- | --- |
| Fixed | Freq. $\geq 0.99$ |
| Majority | Freq. $\geq 0.5$ |
| Minority | $0.01 \leq \text{Freq.} < 0.5$ |

### Variant Annotation

Genetic variants were functionally annotated using SnpEff^77^ (v5.1), based on its official documentation. Annotations included predicted effects on genes and proteins, such as amino acid changes. Genomic feature list of the HIV-1_NL4-3_ strain was obtained from NCBI (accession no. AF324493.2) and was manually updated based on the reconstructed ancestral sequence. The annotation includes nine protein-coding genes and five distinct non-coding regions. Additionally, reversion sites were pinpointed by gene-by-gene alignment of the virus stock’s consensus sequence with the premade 2004 cross-subtype consensus from the Los Alamos HIV Sequence Database. A mutation was classified as a reversion when, in later transfers, the nucleotide at that position changed to match the database consensus.

### TCID_50_ Measurement

We performed titration assay to determine the 50% tissue culture infectious dose (TCID_50_) of evolving HIV-1 populations. For the first 100 transfers, TCID_50_ values were measured for every 10th transfer, and after transfer 100, for every 100 transfers on their respective cell line. Briefly, 1 × 10³ MT-2/4 cells/well were plated in 100 µl of complete medium in two 96-well plates. Serial 1:2.5 dilutions of HIV-1 viral stock were prepared as follows: dilution for row 1 was prepared by mixing 20 µl of cell-free supernatant virus stock with 980 µl of medium (1:50 dilution), followed by a series of 1:3 or 1:4 dilutions (250 µl of the previous dilution mixed with 500/750 µl of medium) until dilution row 11. Row 12 served as the negative control with 500 µl of medium only.

Each viral dilution was added in quadruplicate (50 µl per well) to the pre-seeded MT-2/4 cells. Plates were incubated at 37°C with 5% CO_2_ for 7 days. On day 7 post-infection, viral replication was assessed using microscopic analysis, identifying the presence of syncytia formation and extensive cell debris as indicators of HIV-1 infection. The TCID_50_ values were calculated using the Reed and Muench method^78^, modified by the Spearman-Kaerber formula^79^:

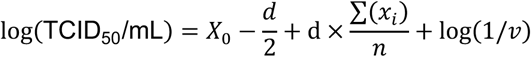

where *X*_0_ represents the initial dilution in the assay, *d* is the log dilution factor, *x_i_* is the number of positive wells per dilution *i*, *n* is the total number of wells, and *v* is the volume of virus inoculum per well.

The multiplicity of infection (MOI) for transfer *t* was then calcualted by the following formula based on a Poisson approximation for the relationship between TCID50 and plaque forming units^80^:

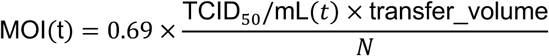

where *N* is the number of seeded cells in each flask (400,000) and transfer_volume is the bottleneck inoculum volume (2 µl).

Number of infectious units per bottleneck at transfer *t* (*IU_bottleneck_*(*t*)) would then be simply yielded from:

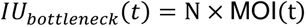

Additionally, to assess cross-infectivity between the two environments, we performed titration assay for viral stock of 100, 200, 300, 400, and 500 transfers of each evolving population on the opposite cell line. The abovementioned formulas were then used to calculate TCID_50_, MOI, and IU*_bottleneck_*(*t*).

### Estimating Effective Population Size

We estimated the effective population size (*N_e_*) under the Wright-Fisher model from the variance in variant frequency change 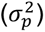 of (nearly) neutral alleles. We used synonymous variants as a simple proxy for (nearly) neutral alleles. 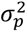 of synonymous variants in the genome after *t* generations was calculated as depicted by the equation below^81^:

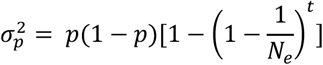

where *p* is the initial variant frequency. If we consider *F* as the standardized variance^82^; where

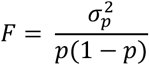

then by approximating for large 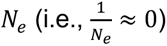, we can derive

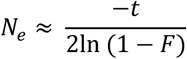

### Genomic RNA Copy Number Quantification

Quantification of HIV-1 genomic RNA copies in cell-free supernatants for every 10th transfer was performed using quantitative PCR (qPCR) with SYBR Green detection on an ABI 7500 real-time PCR system (Applied Biosystems). Reactions were carried out in duplicate in a 20 µl total reaction volume, consisting of 15 µl qPCR master mix and 5 µl of sample (cDNA or standard dilution). The qPCR master mix was prepared using 4 × real-time PCR PRE MIX diluted to 1× final concentration, containing 0.005 µM ROX, 1.5 mM MgCl_2_, 0.4 mM dNTPs, and 0.1× SYBR Green. Specific primers targeting the HIV-1 integrase region (INT:4452-4778) were added at a final concentration of 0.4 µM each, along with JumpStart Taq DNA Polymerase (Sigma-Aldrich) at 1 U per reaction. The qPCR cycling conditions were as follows: initial denaturation at 95°C for 3 min, followed by 50 cycles of denaturation at 94°C for 15 sec, annealing at 55°C for 30 sec, and extension at 72°C for 30 sec. Fluorescence data were collected at the 72°C extension step. A melting curve analysis was conducted immediately after amplification, with a temperature ramp from 55°C to 95°C (1% increase per step), including fluorescence data collection. Negative template controls (NTCs) were included using 5 µl of nuclease-free water in place of the cDNA template. Standard curve quantification was performed using serial dilutions of wild-type (WT) HIV-1 RNA standards to determine absolute RNA copy numbers. The measurements were performed in duplicates for each sample, and the average dilutions were reported.

### Simulation of Neutral Sequence Evolution

To predict how many neutral mutations would arise in our experimental populations, we simulated the evolution of individual HIV-1 genomes of length 9,171 (R-to-R) for 1,500 generations. We started our simulation with a fixed number of 400 infection virions for both MT-2 and MT-4 simulations. Each individual genome was assumed to produce 12 offspring genomes per generation (*R_0_* = 12), and every newly produced viral genome was assumed to acquire a mutation with a probability of 2 × 10^-5^. The mutation target size of neutral positions was assumed to be proportional to the cumulative count of mutated positions at transfer 500. The viral populations were then allowed to replicate exponentially for three generations. Consistent with the experimental setup, we simulated a bottleneck every third generation, by randomly selecting a certain number of individuals from the previous population. The size of the bottleneck, that is, the number of infectious virions transferred at every transfer, was estimated based on infectivity measures experimentally obtained by titration assay for 10th and 100th transfers (refer to “TCID_50_ Measurement” section above). To this end, we estimated IU*_bottleneck_*(*t*) for every transfer by taking the logarithmic average of every 2 available datapoints:

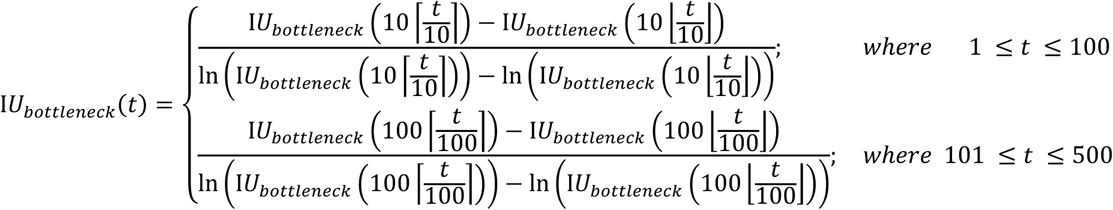

### Parallel Mutation Permutation Tests

#### Neutral parallelism

To estimate the expected level of parallel evolution, we randomly redistributed the fixed mutations observed in each evolutionary line across the genome in the following manner: for each line, fixed mutations were placed at sites where amutation with the same reference and alternative alleles had reached a frequency of at least 0.05 at some point during the experiment. With this approach, we exclude the non-mutated positions (positions which their mutation could results in a lethal effect and hence were not observed) to account for the confounding effects of purifying selection. After random shuffling of fixed mutations over the pool of mutated positions, we measured the proportion of mutations that are shared among each two lines. We then repeated the entire simulation 1,000 times for each pair of lines.

However, this method may still underestimate neutral parallelism, since mutations that occurred later in the experiment might not have had sufficient time to reach fixation. To address this, we subtracted the median fixation time expected under genetic drift (estimated from neutral sequence simulations) from the total experimental duration of 500 transfers and included only mutations that arose before that adjusted time point. Accordingly, we applied median lag corrections of 140 and 185 transfers for the MT-2 and MT-4 lines, respectively.

To determine the expected neutral parallelism for mutations per their translational impact, we performed similar permutation tests. Roughly, 2%, 21%, and 77% of any mutation in the HIV-1 genome would result in untranslated, synonymous, and nonsynonymous mutations, respectively. These percentages were determined based on the consensus ancestral sequence.

#### Environment Contribution

Randomization tests were used to analyze the contribution of environment to observed parallelism within and between each environment as previously described^36^. In short, we redistributed fixed mutations between different experimental lines, while keeping the count of fixed mutations per each line and the number of occurrences of each fixed mutation unchanged. Subsequently, randomized parallelism within and between each environment was calculated and classified as outlined before. The dataset was randomized, and parallelism was classified in this manner for 1,000 times, after which non-parametric significance tests were performed to compare the observed values with the distribution of randomized values.

### Population Diversity Calculation

We calculated and reported Shannon entropy^83^ for population-level genetic diversity. The Shannon index is a measure of diversity that quantifies the information content (entropy) in the sequencing data of a population, representing the uncertainty in predicting the nucleotide base of a locus for a randomly chosen genome. Shannon indices were obtained according to the following formula for each locus:

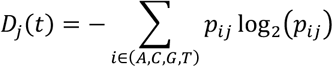

where *D_j_*(*t*) is entropy at locus *j* at transfer *t*, and *p_ij_* is the proportion of each of the 4 possible nucleotides (A, C, G, T). We obtained the mean diversity 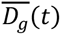 over the HIV-1 genome with length *L_g_*:

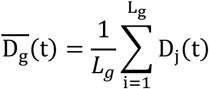

### dN/dS Calculation

dN and dS were calculated and reported according to the following definitions:

- dN(*t*) = number of nonsynonymous mutations at transfer *t* / total number of nonsynonymous sites
- dS(*t*) = number of synonymous substitutions at transfer *t* / total number of synonymous sites

### Time to fixation measurement

To estimate the time to fixation of mutations, we tracked their frequencies across passages for each experimental line. For each line, mutations that reached fixation (frequency ≥ 0.99 at transfer 500) were identified. The transfer at which a mutation first appeared was determined as the earliest passage with frequency ≥ 0.01, and the passage of fixation was recorded as the earliest passage with frequency ≥ 0.99. If the frequency after appearance dropped to below 0.01 for two consecutive transfers, the transfer of appearance was updated to the next transfer it reappeared. With this method, we computed the continuous time it took a mutation to appear and reach fixation.

### Linkage disequilibrium

Linkage disequilibrium (LD) between single-base mutations was calculated using paired-end sequencing data with an in-house script. Mutations with a frequency ≥ 0.05 in each transfer were considered. For each pair of mutations separated by ≤ 400 bp, haplotypes were reconstructed from read pairs covering both positions. Reads and their mates were aligned to the reference genome, and nucleotides at the mutation positions were extracted to define haplotypes. Frequencies of the four possible haplotypes (reference-reference (*fab*), reference-alternate (*faB*), alternate-reference (*fAb*), alternate-alternate (*fAB*)) were obtained. The disequilibrium coefficient (D) was calculated as follow:

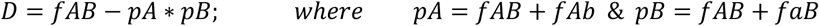

and the squared correlation coefficient (*r*^2^) as a measure of LD was subsequently calculated as:

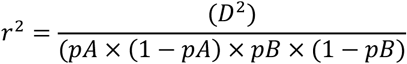

Mutation pairs with no overlapping reads were assigned NA. LD calculations were performed in parallel across available cores to optimize computational speed.

### Initial relative fitness calculation

To assess the relationship between fitness and fixation probability, we estimated initial relative fitness as described in Lang *et al.* (2011). In short, for each mutation, we identified *t*_1_ and *t*_2_, the first consecutive transfers where the frequency of the mutation was greater than 0.01 and 0.1, respectively. We then calculated initial relative fitness as:

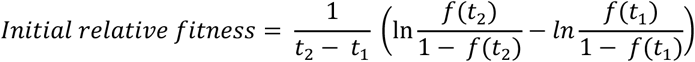

where f(*t*) is the frequency of the mutation at transfer *t*.

### Website

The interactive web application for exploring and accessing the curated data was developed using the R Shiny package and is available at: https://lteeviz.shinyapps.io/lteeviz/

### Use of Artificial Intelligence

We acknowledge the use of OpenAI’s ChatGPT (GPT-4 and GPT-5) for language edits and Anthropic’s Claude AI to assist with code debugging and syntax refinement. All text and code logic, design, and interpretation were performed and verified by the authors.

## Code and data availability

All the code for data processing pipeline and downstream analyses can be found on GitHub at git@github.com:Al313/hiv1_genome_evolution.git. The Illumina sequencing data were deposited in the NCBI short read archive under the accession number PRJNA506879. The code for web application tool (LTEEviz) can be found on GitHub at git@github.com:Al313/lte-viz.git. The curated data can be obtained via LTEEviz.

## Supplementary Material

Supplementary data are enclosed in a single PDF and will be made available online.

## Acknowledgement

C.L., K.N., R.C. and K.J.M. performed the experiments; K.J.M. and R.R.R. designed the experiments; A.M. analyzed the data; A.M. wrote the article; R.R.R. and K.J.M. provided feedback on the article; A.M. designed, coded, and implemented LTEEviz web application tool; L.S. contributed to the coding of LTEEviz. A.M. and L.S. were funded by European Union’s Horizon 2020 research and innovation program, under the Marie Skłodowska-Curie Actions Innovative Training Networks grant no. 955974 (VIROINF). K.J.M. acknowledges the financial support of the Swiss National Science Foundation (grant no. SNF 204404). K.J.M. acknowledges the financial support of the 2021 fellowship program by Gilead Sciences (grant no. 13849).

## Reference

1. Silander, O. K., Tenaillon, O. & Chao, L. Understanding the Evolutionary Fate of Finite Populations: The Dynamics of Mutational Effects. PLOS Biol. 5, e94 (2007).

2. Wiser, M. J., Ribeck, N. & Lenski, R. E. Long-Term Dynamics of Adaptation in Asexual Populations. Science 342, 1364–1367 (2013).

3. Kryazhimskiy, S., Rice, D. P., Jerison, E. R. & Desai, M. M. Global epistasis makes adaptation predictable despite sequence-level stochasticity. Science 344, 1519–1522 (2014).

4. Losos, J. B., Jackman, T. R., Larson, A., Queiroz, K. de & Rodriguez-Schettino, L. Contingency and Determinism in Replicated Adaptive Radiations of Island Lizards. Science 279, 2115–2118 (1998).

5. Chou, H.-H., Chiu, H.-C., Delaney, N. F., Segrè, D. & Marx, C. J. Diminishing Returns Epistasis Among Beneficial Mutations Decelerates Adaptation. Science 332, 1190–1192 (2011).

6. Blount, Z. D., Lenski, R. E. & Losos, J. B. Contingency and determinism in evolution: Replaying life’s tape. Science 362, eaam5979 (2018).

7. Lenormand, T., Roze, D. & Rousset, F. Stochasticity in evolution. Trends Ecol. Evol. 24, 157–165 (2009).

8. Schoustra, S. E., Punzalan, D., Dali, R., Rundle, H. D. & Kassen, R. Multivariate Phenotypic Divergence Due to the Fixation of Beneficial Mutations in Experimentally Evolved Lineages of a Filamentous Fungus. PLOS ONE 7, e50305 (2012).

9. Jensen, J. D. & Bachtrog, D. Characterizing the Influence of Effective Population Size on the Rate of Adaptation: Gillespie’s Darwin Domain. Genome Biol. Evol. 3, 687–701 (2011).

10. Papkou, A., Garcia-Pastor, L., Escudero, J. A. & Wagner, A. A rugged yet easily navigable fitness landscape. Science 382, eadh3860 (2023).

11. Vitti, J. J., Grossman, S. R. & Sabeti, P. C. Detecting Natural Selection in Genomic Data. Annu. Rev. Genet. 47, 97–120 (2013).

12. Gillespie, John H. The Causes of Molecular Evolution. (Oxford University Press, 1992).

13. Barrett, R. D. H. & Schluter, D. Adaptation from standing genetic variation. Trends Ecol. Evol. 23, 38–44 (2008).

14. Couce, A. & Tenaillon, O. A. The rule of declining adaptability in microbial evolution experiments. Front. Genet. 6, (2015).

15. Wünsche, A. et al. Diminishing-returns epistasis decreases adaptability along an evolutionary trajectory. *Nat*. Ecol. Evol. 1, 0061 (2017).

16. Booker, T. R., Jackson, B. C. & Keightley, P. D. Detecting positive selection in the genome. BMC Biol. 15, 98 (2017).

17. Orr, H. A. The genetic theory of adaptation: a brief history. Nat. Rev. Genet. 6, 119–127 (2005).

18. Lenski, R. E. Experimental evolution and the dynamics of adaptation and genome evolution in microbial populations. ISME J. 11, 2181–2194 (2017).

19. Foote, M. & Raup, D. M. Fossil preservation and the stratigraphic ranges of taxa. Paleobiology 22, 121–140 (1996).

20. Benson, R. B. J., Butler, R., Close, R. A., Saupe, E. & Rabosky, D. L. Biodiversity across space and time in the fossil record. Curr. Biol. 31, R1225–R1236 (2021).

21. Trappes, R. Defining the niche for niche construction: evolutionary and ecological niches. Biol. Philos. 36, 31 (2021).

22. Lenski, R. E., Rose, M. R., Simpson, S. C. & Tadler, S. C. Long-Term Experimental Evolution in Escherichia coli. I. Adaptation and Divergence During 2,000 Generations. Am. Nat. 138, 1315–1341 (1991).

23. Wichman, H. A., Scott, L. A., Yarber, C. D. & Bull, J. J. Experimental evolution recapitulates natural evolution. Philos. Trans. R. Soc. B Biol. Sci. 355, 1677–1684 (2000).

24. Burke, M. K. et al. Genome-wide analysis of a long-term evolution experiment with Drosophila. Nature 467, 587–590 (2010).

25. Lang, G. I. et al. Pervasive genetic hitchhiking and clonal interference in forty evolving yeast populations. Nature 500, 571–574 (2013).

26. Tenaillon, O. et al. Tempo and mode of genome evolution in a 50,000-generation experiment. Nature 536, 165–170 (2016).

27. Feder, A. F. et al. A spatio-temporal assessment of simian/human immunodeficiency virus (SHIV) evolution reveals a highly dynamic process within the host. PLoS Pathog. 13, e1006358 (2017).

28. Carr, A., Mackie, N. E., Paredes, R. & Ruxrungtham, K. HIV drug resistance in the era of contemporary antiretroviral therapy: A clinical perspective. Antivir. Ther. 28, 13596535231201162 (2023).

29. Mansky, L. M. & Temin, H. M. Lower in vivo mutation rate of human immunodeficiency virus type 1 than that predicted from the fidelity of purified reverse transcriptase. J. Virol. 69, 5087–5094 (1995).

30. Kouyos, R. D., Althaus, C. L. & Bonhoeffzer, S. Stochastic or deterministic: what is the effective population size of HIV-1? Trends Microbiol. 14, 507–511 (2006).

31. Rodrigo, A. G. et al. Coalescent estimates of HIV-1 generation time in vivo. Proc. Natl. Acad. Sci. U. S. A. 96, 2187–2191 (1999).

32. Bertels, F., Leemann, C., Metzner, K. J. & Regoes, R. R. Parallel Evolution of HIV-1 in a Long-Term Experiment. Mol. Biol. Evol. 36, 2400–2414 (2019).

33. Woods, R., Schneider, D., Winkworth, C. L., Riley, M. A. & Lenski, R. E. Tests of parallel molecular evolution in a long-term experiment with Escherichia coli. Proc. Natl. Acad. Sci. 103, 9107–9112 (2006).

34. Burke, M. K. et al. Genome-wide analysis of a long-term evolution experiment with Drosophila. Nature 467, 587–590 (2010).

35. Kvitek, D. J. & Sherlock, G. Whole Genome, Whole Population Sequencing Reveals That Loss of Signaling Networks Is the Major Adaptive Strategy in a Constant Environment. PLOS Genet. 9, e1003972 (2013).

36. Bons, E., Leemann, C., Metzner, K. J. & Regoes, R. R. Long-term experimental evolution of HIV-1 reveals effects of environment and mutational history. PLOS Biol. 18, e3001010 (2020).

37. Ribeiro, R. M. et al. Estimation of the Initial Viral Growth Rate and Basic Reproductive Number during Acute HIV-1 Infection. J. Virol. 84, 6096–6102 (2010).

38. Iwami, S. et al. Cell-to-cell infection by HIV contributes over half of virus infection. eLife 4, e08150 (2015).

39. Mohammadi, P. et al. 24 Hours in the Life of HIV-1 in a T Cell Line. PLOS Pathog. 9, e1003161 (2013).

40. Perelson, A. HIV-1 Dynamics in Vivo: Virion Clearance Rate, Infected Cell Life-Span, and Viral Generation Time. Am. Assoc. Adv. Sci. 10.1126/science.271.5255.1582 (1996) doi:10.1126/science.271.5255.1582.

41. Crow, J. & Kimura, M. An Introduction to Population Genetics Theory. (Blackburn Press, 1970).

42. Charlesworth, B. & Charlesworth, D. Elements of Evolutionary Genetics. (Roberts & Company Publishers, 2010).

43. Abram, M. E., Ferris, A. L., Shao, W., Alvord, W. G. & Hughes, S. H. Nature, Position, and Frequency of Mutations Made in a Single Cycle of HIV-1 Replication. J. Virol. 84, 9864–9878 (2010).

44. Abram, M. E. et al. Mutations in HIV-1 Reverse Transcriptase Affect the Errors Made in a Single Cycle of Viral Replication. J. Virol. 88, 7589–7601 (2014).

45. Rawson, J. M. O., Landman, S. R., Reilly, C. S. & Mansky, L. M. HIV-1 and HIV-2 exhibit similar mutation frequencies and spectra in the absence of G-to-A hypermutation. Retrovirology 12, 60 (2015).

46. Shankarappa, R. et al. Consistent Viral Evolutionary Changes Associated with the Progression of Human Immunodeficiency Virus Type 1 Infection. J. Virol. 73, 10489–10502 (1999).

47. Zanini, F. et al. Population genomics of intrapatient HIV-1 evolution. eLife 4, e11282 (2015).

48. Wichman, H. A., Badgett, M. R., Scott, L. A., Boulianne, C. M. & Bull, J. J. Different trajectories of parallel evolution during viral adaptation. Science 285, 422–424 (1999).

49. Grantham, R. Amino Acid Difference Formula to Help Explain Protein Evolution. Science 185, 862–864 (1974).

50. Gillespie, J. H. Genetic Drift in an Infinite Population: The Pseudohitchhiking Model. Genetics 155, 909–919 (2000).

51. DePristo, M. A., Weinreich, D. M. & Hartl, D. L. Missense meanderings in sequence space: a biophysical view of protein evolution. Nat. Rev. Genet. 6, 678–687 (2005).

52. Zhang, J., Chen, F. & Li, X. Mechanistic causes of sign epistasis and its applications. Front. Genet. 15, (2024).

53. Lang, G. I., Botstein, D. & Desai, M. M. Genetic Variation and the Fate of Beneficial Mutations in Asexual Populations. Genetics 188, 647–661 (2011).

54. Smith, J. M. & Haigh, J. The hitch-hiking effect of a favourable gene. Genet. Res. 23, 23–35 (1974).

55. Zhao, S., Chi, L., Fu, M. & Chen, H. HaploSweep: Detecting and Distinguishing Recent Soft and Hard Selective Sweeps through Haplotype Structure. Mol. Biol. Evol. 41, msae192 (2024).

56. Rokyta, D. R., Abdo, Z. & Wichman, H. A. The Genetics of Adaptation for Eight Microvirid Bacteriophages. J. Mol. Evol. 69, 229–239 (2009).

57. Schoustra, S. E., Bataillon, T., Gifford, D. R. & Kassen, R. The Properties of Adaptive Walks in Evolving Populations of Fungus. PLOS Biol. 7, e1000250 (2009).

58. Neidhart, J. & Krug, J. Adaptive Walks and Extreme Value Theory. Phys. Rev. Lett. 107, 178102 (2011).

59. Wielgoss, S. et al. Mutation Rate Inferred From Synonymous Substitutions in a Long-Term Evolution Experiment With Escherichia coli. G3 GenesGenomesGenetics 1, 183–186 (2011).

60. Kim, Y. & Orr, H. A. Adaptation in Sexuals vs. Asexuals: Clonal Interference and the Fisher-Muller Model. Genetics 171, 1377–1386 (2005).

61. Muller, H. J. Some Genetic Aspects of Sex. Am. Nat. 66, 118–138 (1932).

62. Fisher, R. A. The Genetical Theory of Natural Selection. (Oxford University Press, Oxford, New York, 1930).

63. Goddard, M. R., Godfray, H. C. J. & Burt, A. Sex increases the efficacy of natural selection in experimental yeast populations. Nature 434, 636–640 (2005).

64. Maddamsetti, R., Lenski, R. E. & Barrick, J. E. Adaptation, Clonal Interference, and Frequency-Dependent Interactions in a Long-Term Evolution Experiment with Escherichia coli. Genetics 200, 619–631 (2015).

65. Simon-Loriere, E. & Holmes, E. C. Why do RNA viruses recombine? Nat. Rev. Microbiol. 9, 617–626 (2011).

66. Moradigaravand, D. et al. Recombination Accelerates Adaptation on a Large-Scale Empirical Fitness Landscape in HIV-1. PLOS Genet. 10, e1004439 (2014).

67. Wichman, H. A., Millstein, J. & Bull, J. J. Adaptive Molecular Evolution for 13,000 Phage Generations: A Possible Arms Race. Genetics 170, 19–31 (2005).

68. Bauer, B. & Gokhale, C. S. Repeatability of evolution on epistatic landscapes. Sci. Rep. 5, 9607 (2015).

69. Sackman, A. M. et al. Mutation-Driven Parallel Evolution during Viral Adaptation. Mol. Biol. Evol. 34, 3243–3253 (2017).

70. Lind, P. A., Libby, E., Herzog, J. & Rainey, P. B. Predicting mutational routes to new adaptive phenotypes. eLife 8, e38822 (2019).

71. Svensson, E. I. & Berger, D. The Role of Mutation Bias in Adaptive Evolution. Trends Ecol. Evol. 34, 422–434 (2019).

72. Bohn, P., Gribling-Burrer, A.-S., Ambi, U. B. & Smyth, R. P. Nano-DMS-MaP allows isoform-specific RNA structure determination. Nat. Methods 20, 849–859 (2023).

73. Sarashetti, P., Lipovac, J., Tomas, F., Šikić, M. & Liu, J. Evaluating data requirements for high-quality haplotype-resolved genomes for creating robust pangenome references. Genome Biol. 25, 312 (2024).

74. Giallonardo, F. D. et al. Full-length haplotype reconstruction to infer the structure of heterogeneous virus populations. Nucleic Acids Res. 42, e115 (2014).

75. Harada, S., Koyanagi, Y. & Yamamoto, N. Infection of HTLV-III/LAV in HTLV-I-Carrying Cells MT-2 and MT-4 and Application in a Plaque Assay. Science 229, 563–566 (1985).

76. Adachi, A. et al. Production of acquired immunodeficiency syndrome-associated retrovirus in human and nonhuman cells transfected with an infectious molecular clone. J. Virol. 59, 284–291 (1986).

77. Cingolani, P. et al. A program for annotating and predicting the effects of single nucleotide polymorphisms, SnpEff: SNPs in the genome of Drosophila melanogaster strain w1118; iso-2; iso-3. Fly (Austin) 6, 80–92 (2012).

78. Reed, L. J. & Muench, H. A SIMPLE METHOD OF ESTIMATING FIFTY PER CENT ENDPOINTS12. Am. J. Epidemiol. 27, 493–497 (1938).

79. Kärber, G. Beitrag zur kollektiven Behandlung pharmakologischer Reihenversuche. Naunyn-Schmiedebergs Arch. Für Exp. Pathol. Pharmakol. 162, 480–483 (1931).

80. Bryan, W. R. Interpretation of Host Response in Quantitative Studies on Animal Viruses. Ann. N. Y. Acad. Sci. 69, 698–728 (1957).

81. Jónás, Á., Taus, T., Kosiol, C., Schlötterer, C. & Futschik, A. Estimating the Effective Population Size from Temporal Allele Frequency Changes in Experimental Evolution. Genetics 204, 723–735 (2016).

82. Wright, Swell. Evolution in Mendelian Populations. (Genetics, 1931).

83. Sherwin, W. B. Entropy and Information Approaches to Genetic Diversity and its Expression: Genomic Geography. Entropy 12, 1765–1798 (2010).

