## Supplementary Material for "Substantial Deceleration of Adaptation of HIV-1 Within 1,500 Generations in an Experimental Evolution: A Genomic Perspective"

**This PDF file includes**

Supplementary Figs. S1 – S17

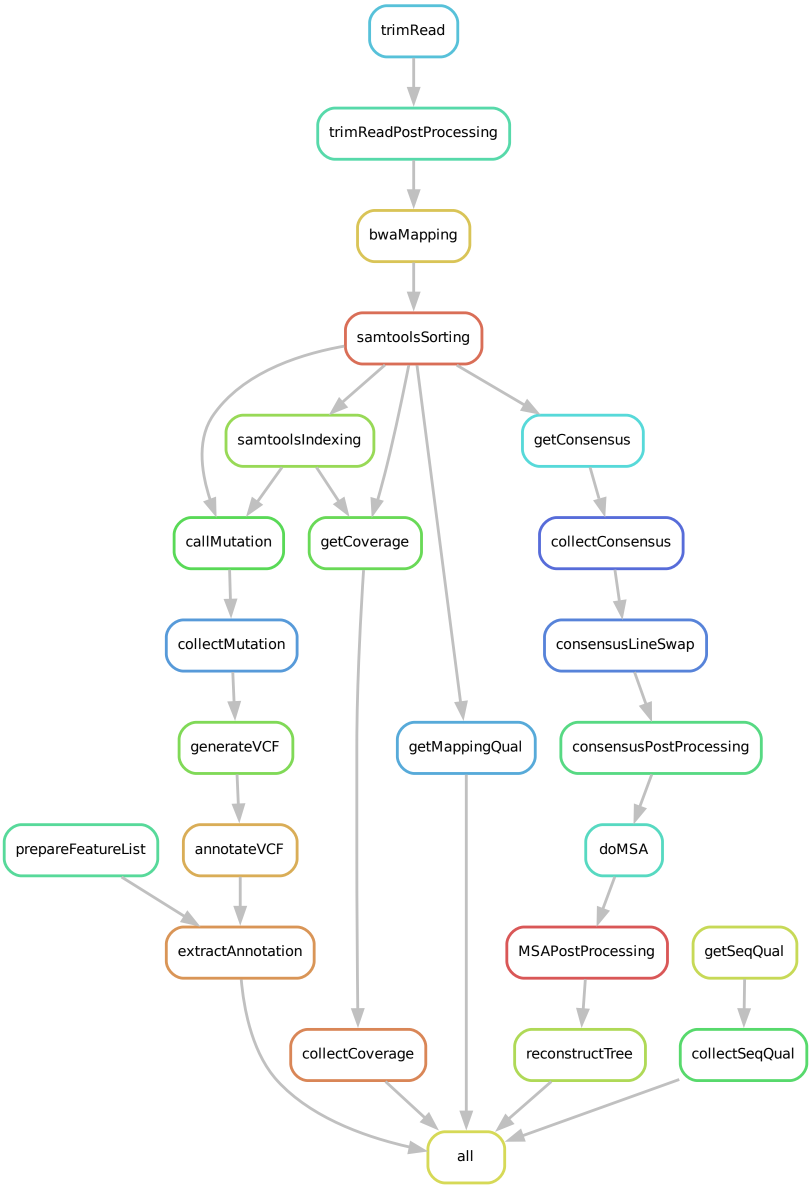


**Fig. S1. Data processing pipeline.** All the data processing was performed via a single Snakemake pipeline. The pipeline starts from raw sequencing reads per sample and ends with the rule “all”, requiring as output annotated variants, sequencing and mapping quality, genome read coverage, and reconstructed phylogeny tree.

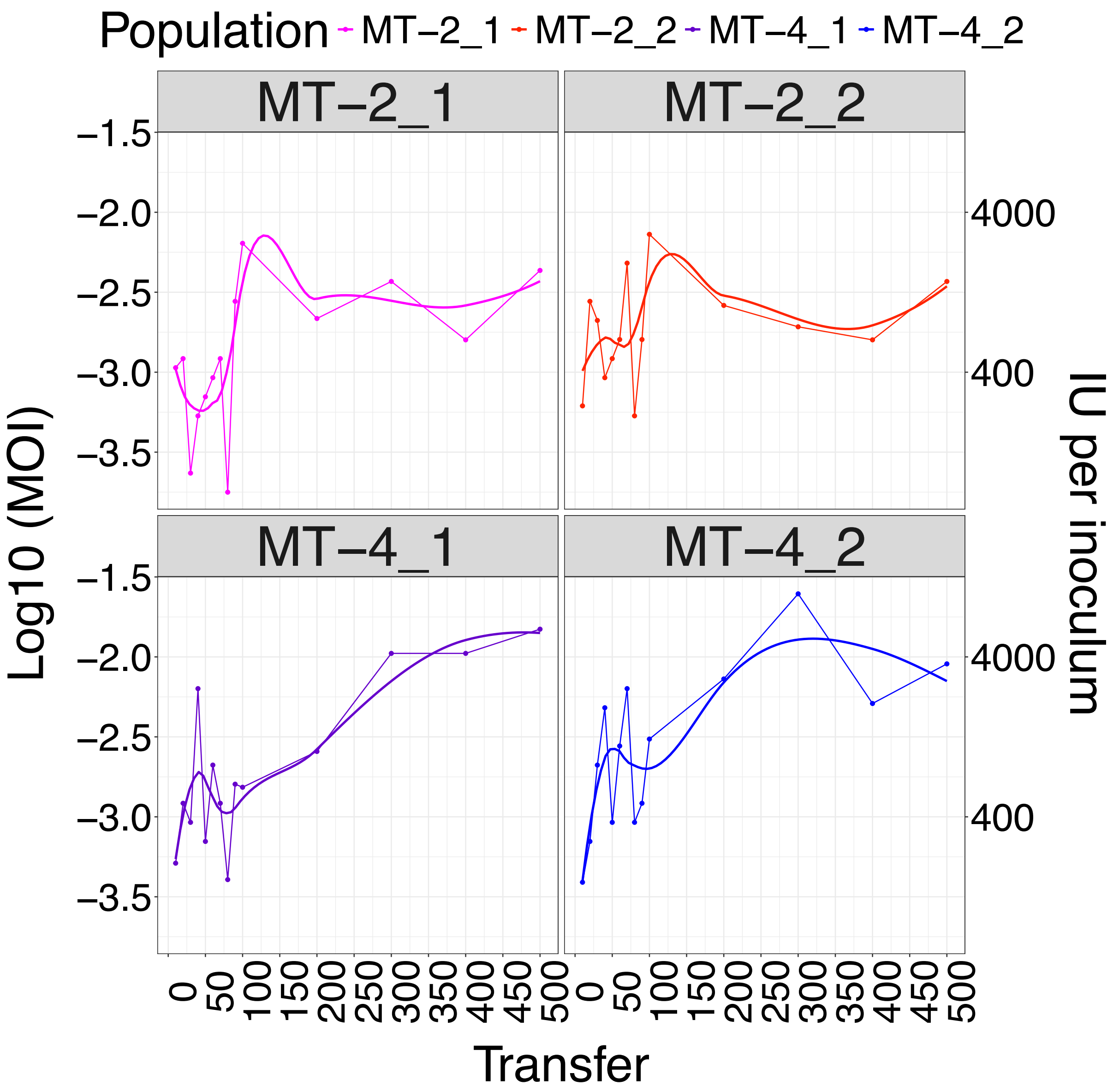


**Fig. S2. Multiplicity of infection data.** Titration assay was performed for every 10th transfer until transfer 100, and for every 100th transfer until transfer 500. The left axis shows the MOI on log10 base, and the right axis shows number of infectious units per 2 µl of transferred cell suspension at every transfer.

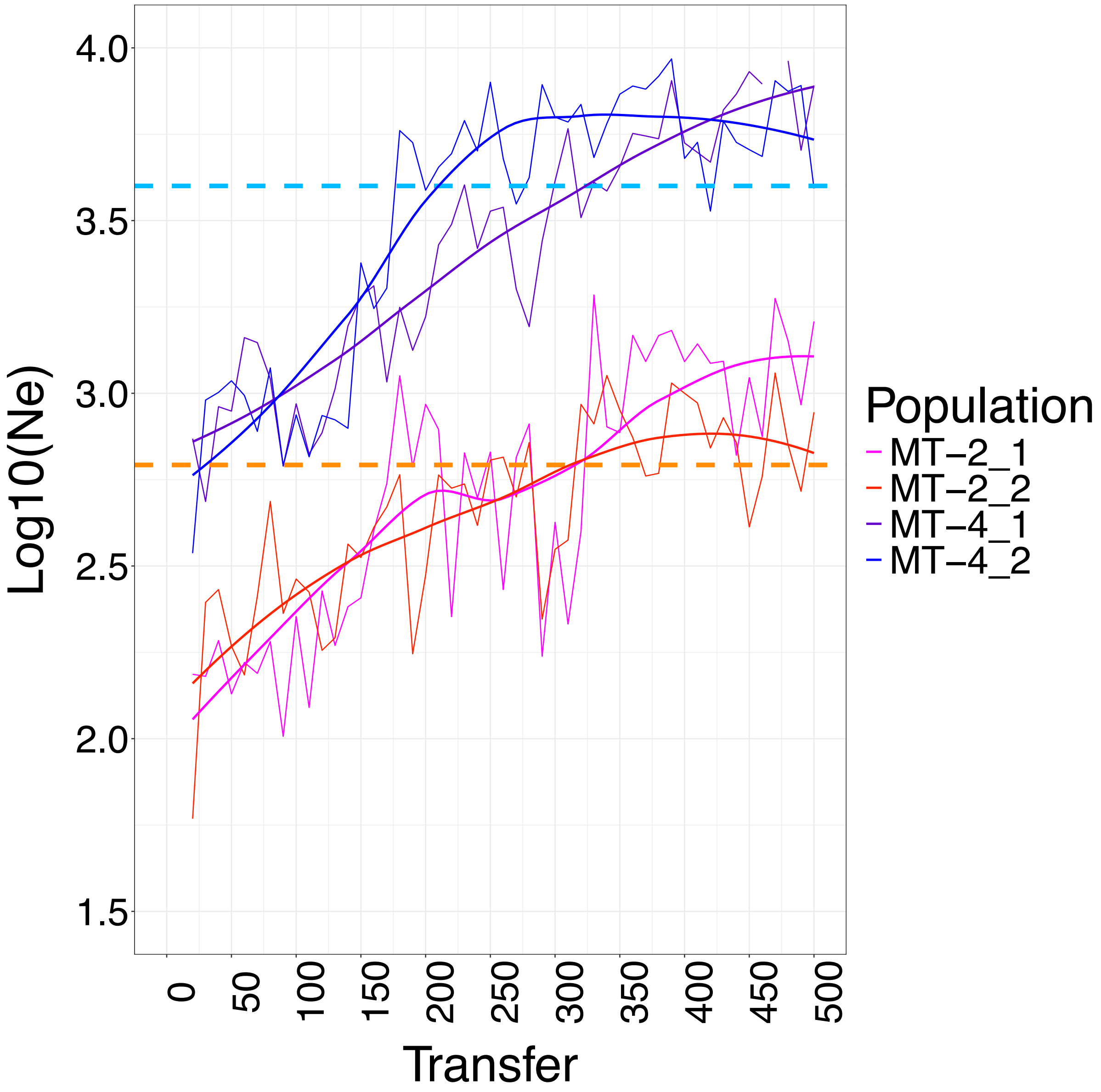


**Fig. S3. Variance effective population size.** Estimates of effective population size ($N_e$) based on average variance in frequency of mutations in subsequent samples. The orange and blue horizontal dashed lines show the mean $N_e$ for MT-2 (2.8) and MT-4 (3.6) replicates over time, respectively.

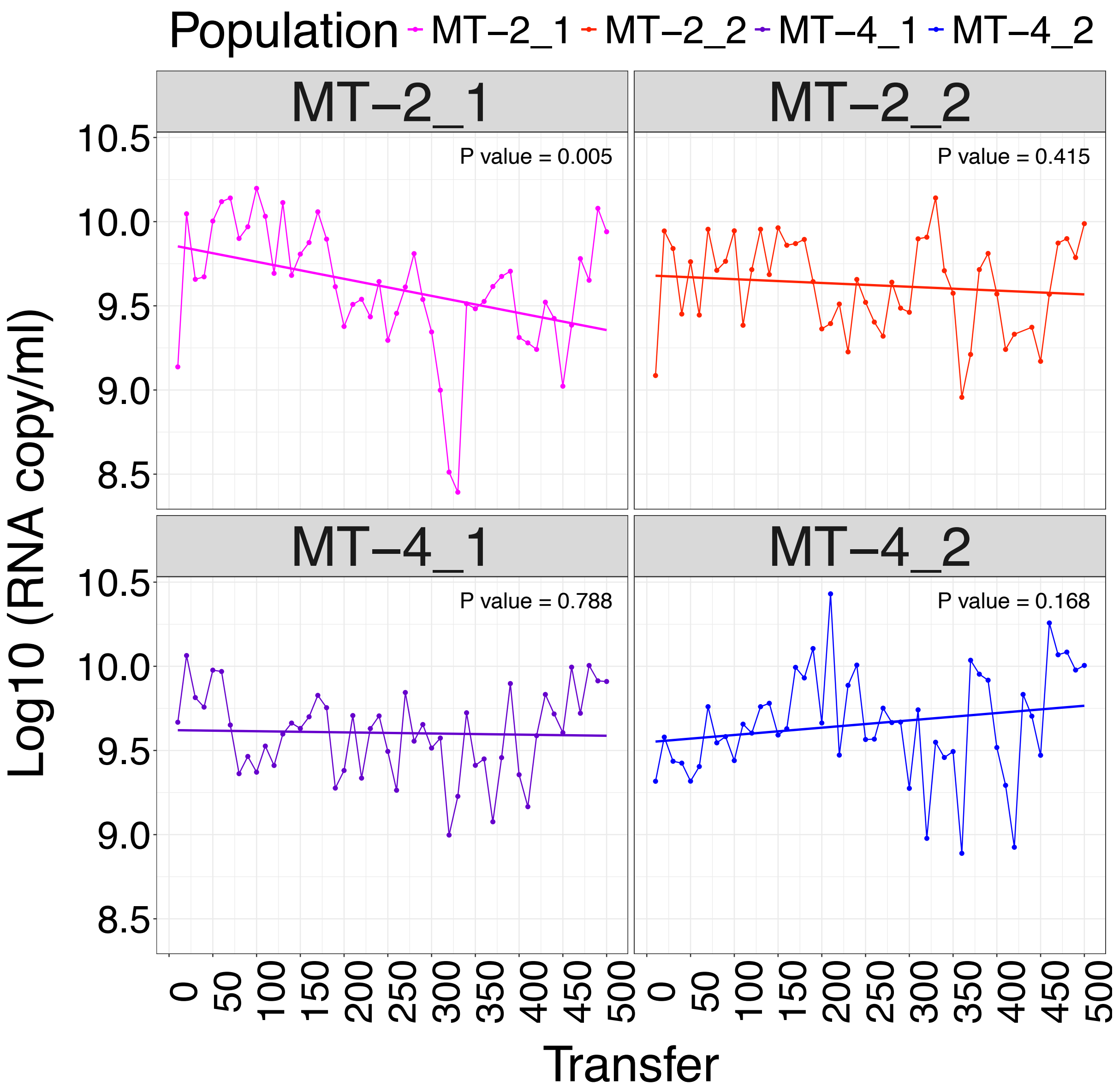


**Fig. S4. Kinetics of viral titers.** Quantitative PCR was performed for every 10th transfer until transfer 500. The qPCR is expected to measure the RNA copy number in cell suspension supernatants (both genomic and transcriptomic RNA). Linear regression did not result in a significant up/down trend for all evolution lines, but MT-2_1.

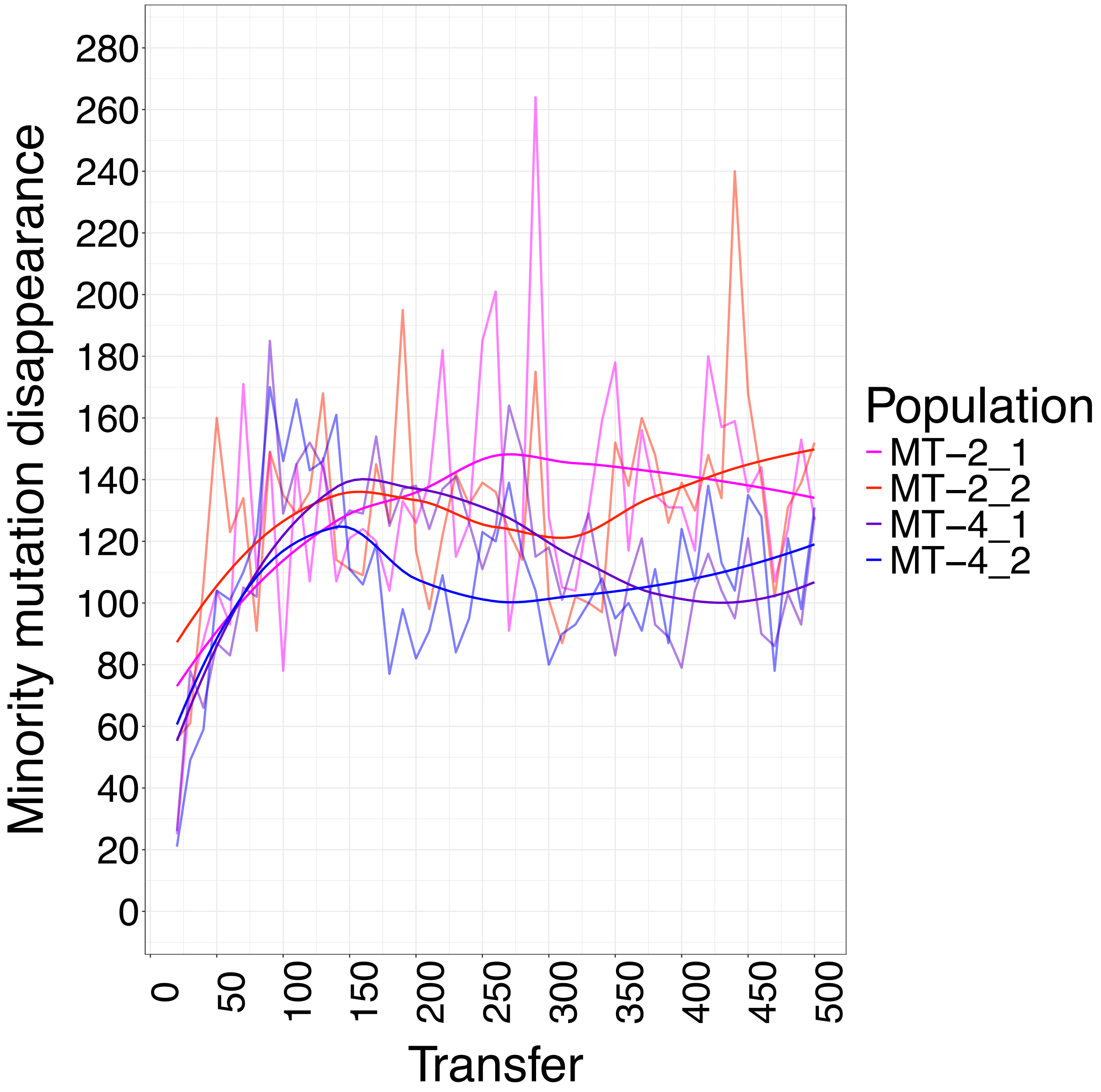


**Fig. S5. Rate of minority mutation disappearance.** Number of minority mutations that were present at transfer *t* but not in transfer *t+10*. Rate of minority mutation was significant higher in MT-2 replicates than MT-4 ($P < 0.001$, paired t-test).

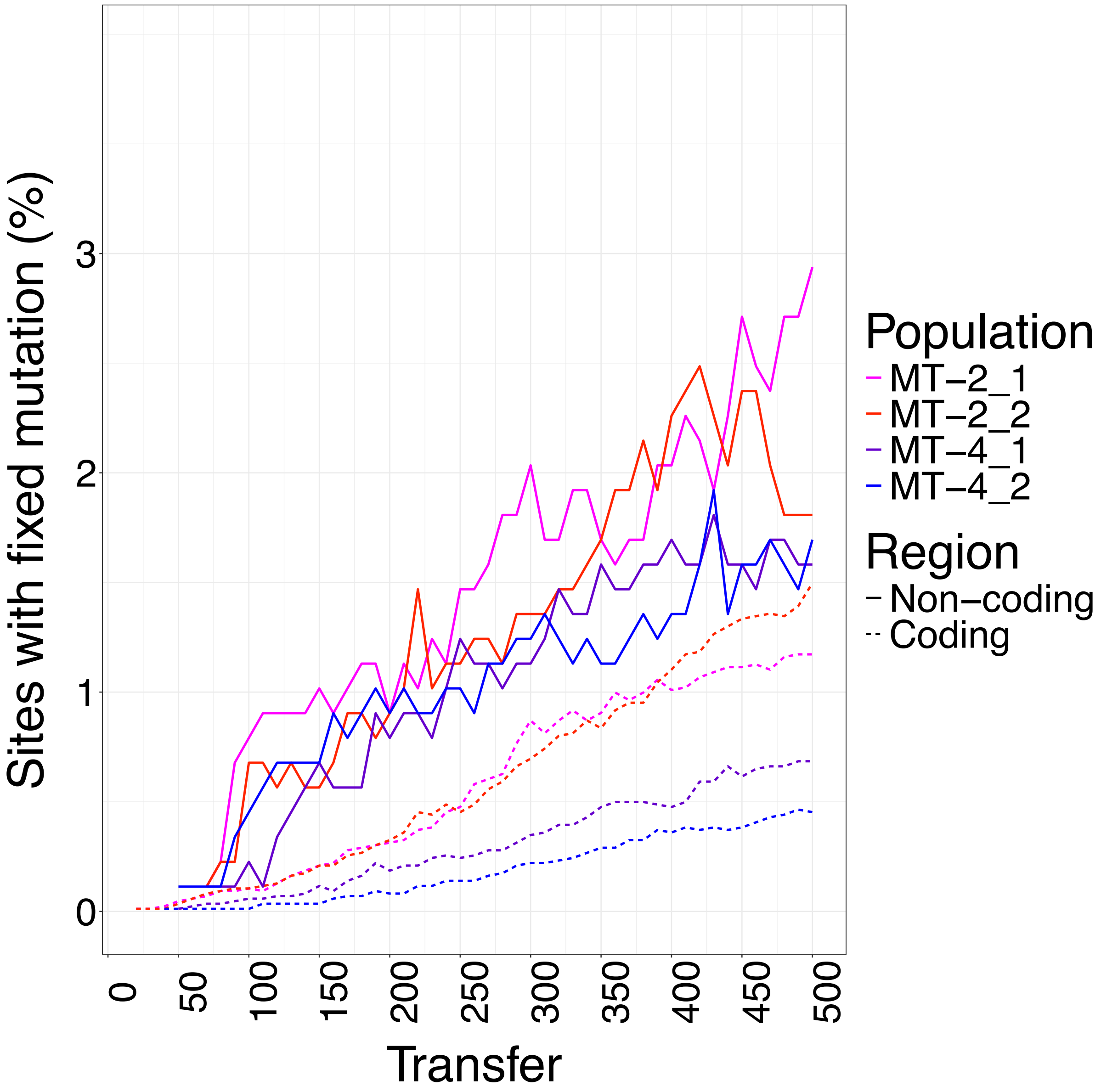


**Fig. S6. Fixed mutation accumulation patterns per coding and non-coding regions.** Fraction of coding and non-coding sites with a fixed mutation.

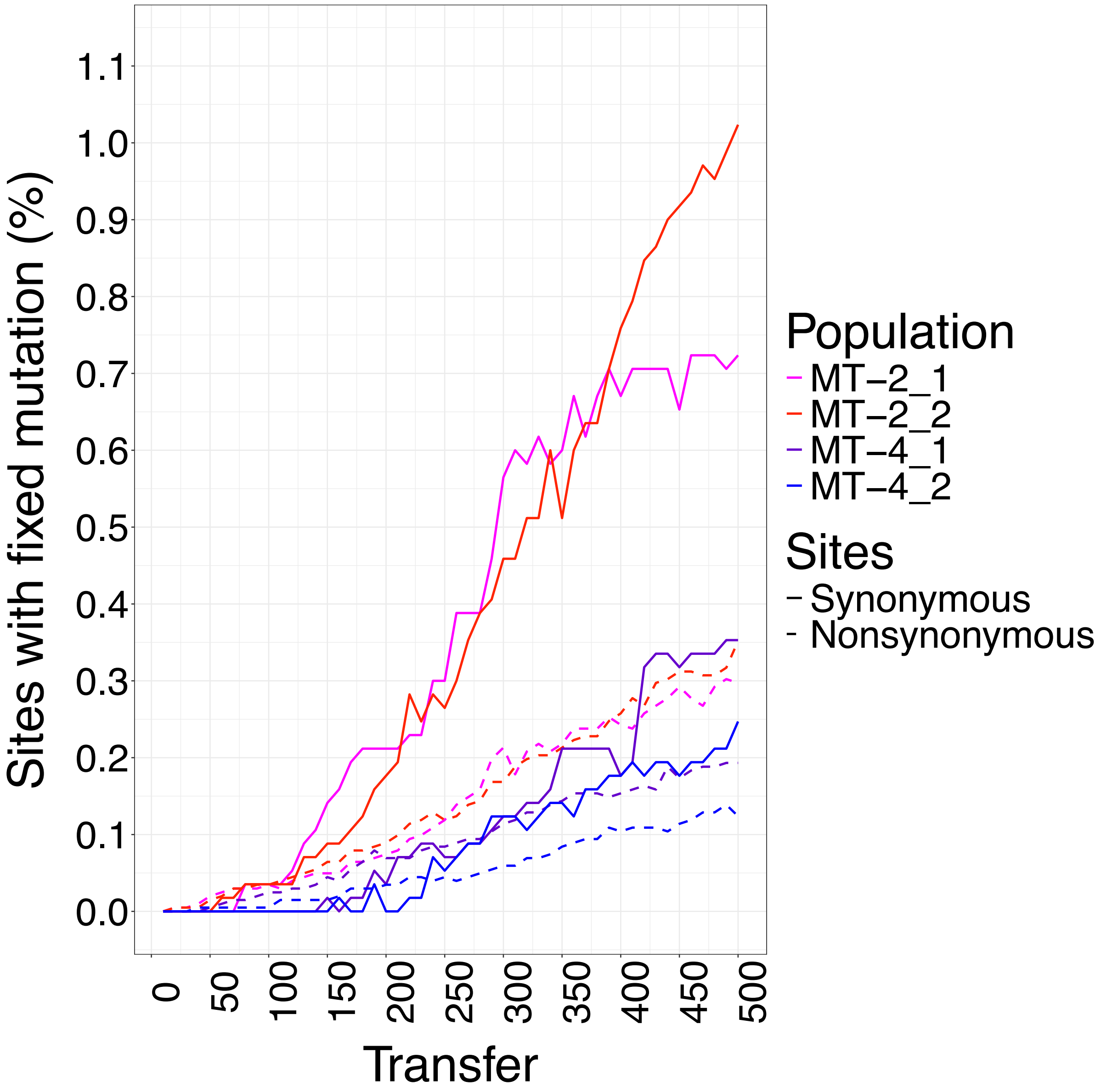


**Fig. S7. Fixed mutation accumulation patterns per synonymous and nonsynonymous sites.**

Fraction of synonymous and nonsynonymous sites with a fixed mutation.

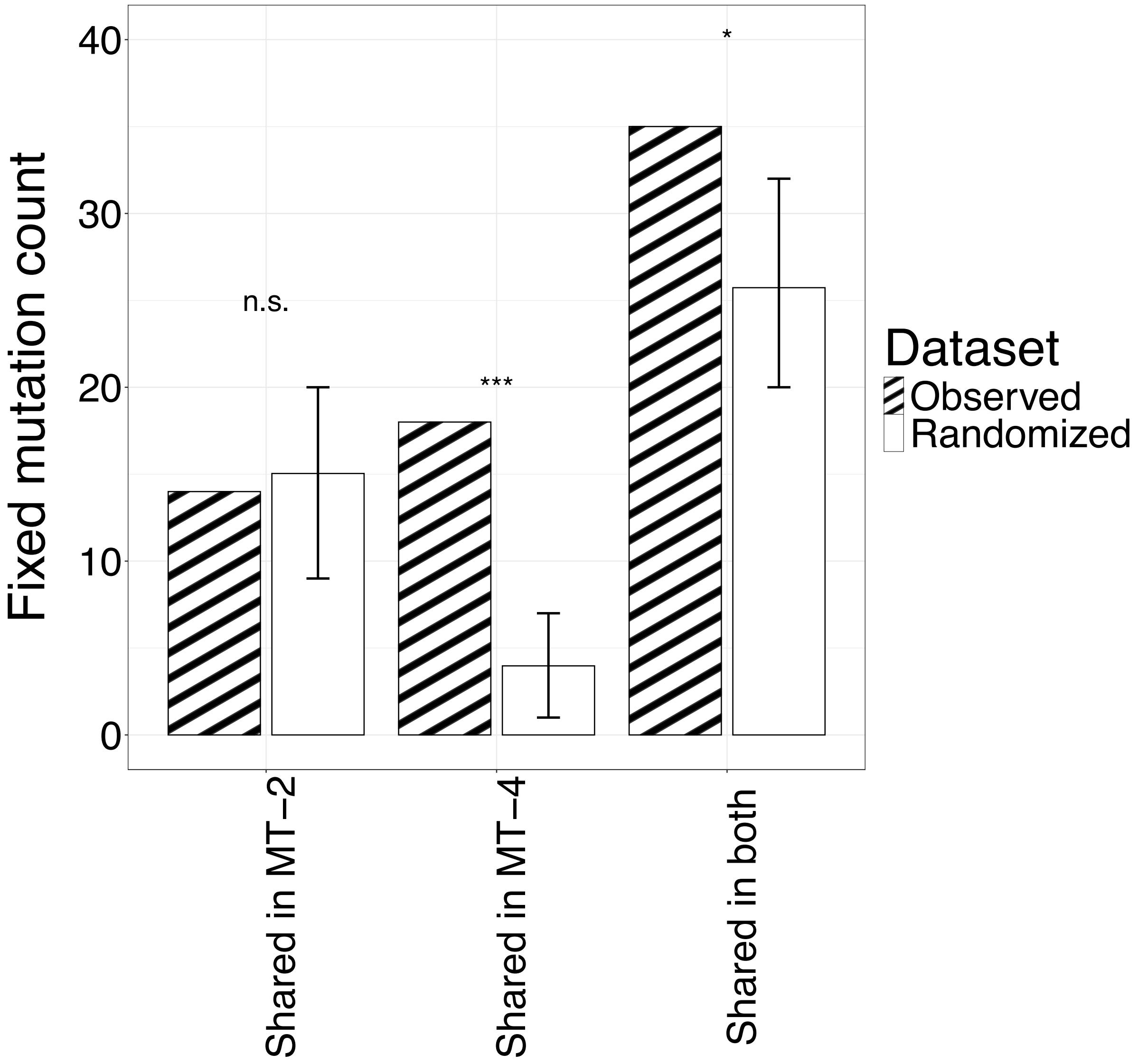


**Fig. S8. Environment randomization tests for expected parallelism.** Level of observed and expected parallelism among fixed mutations. Randomization test was performed by redistributing fixed mutations between different experimental lines, while keeping the count of fixed mutations per each line and the number of occurrences of each fixed mutation unchanged. Parallelism among MT-4 replicates was significantly different than expected, whereas for MT-2 replicates the difference was not significant.

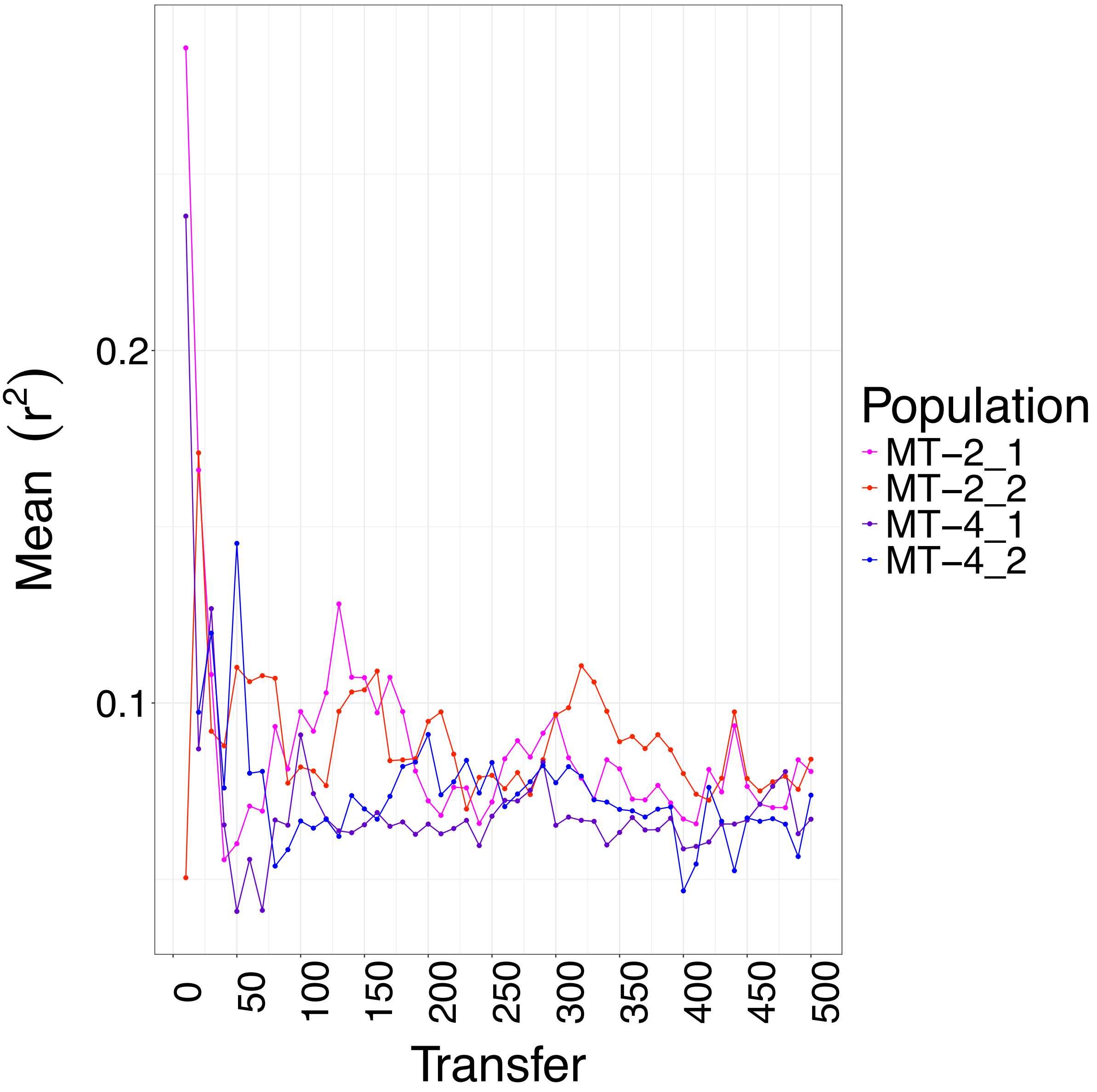


**Fig. S9. Linkage disequilibrium per sample**. We measured linkage disequilibrium coefficient for mutations that were covered by the same read pair (maximum distance of 400 bp, see Methods). We then calculated squared correlation coefficient of alleles ($r^2$) for these mutations and averaged them over the length of genome. In early samples, $r^2$ values showed greater variation, likely due to the smaller number of mutation pairs contributing to the final calculation.

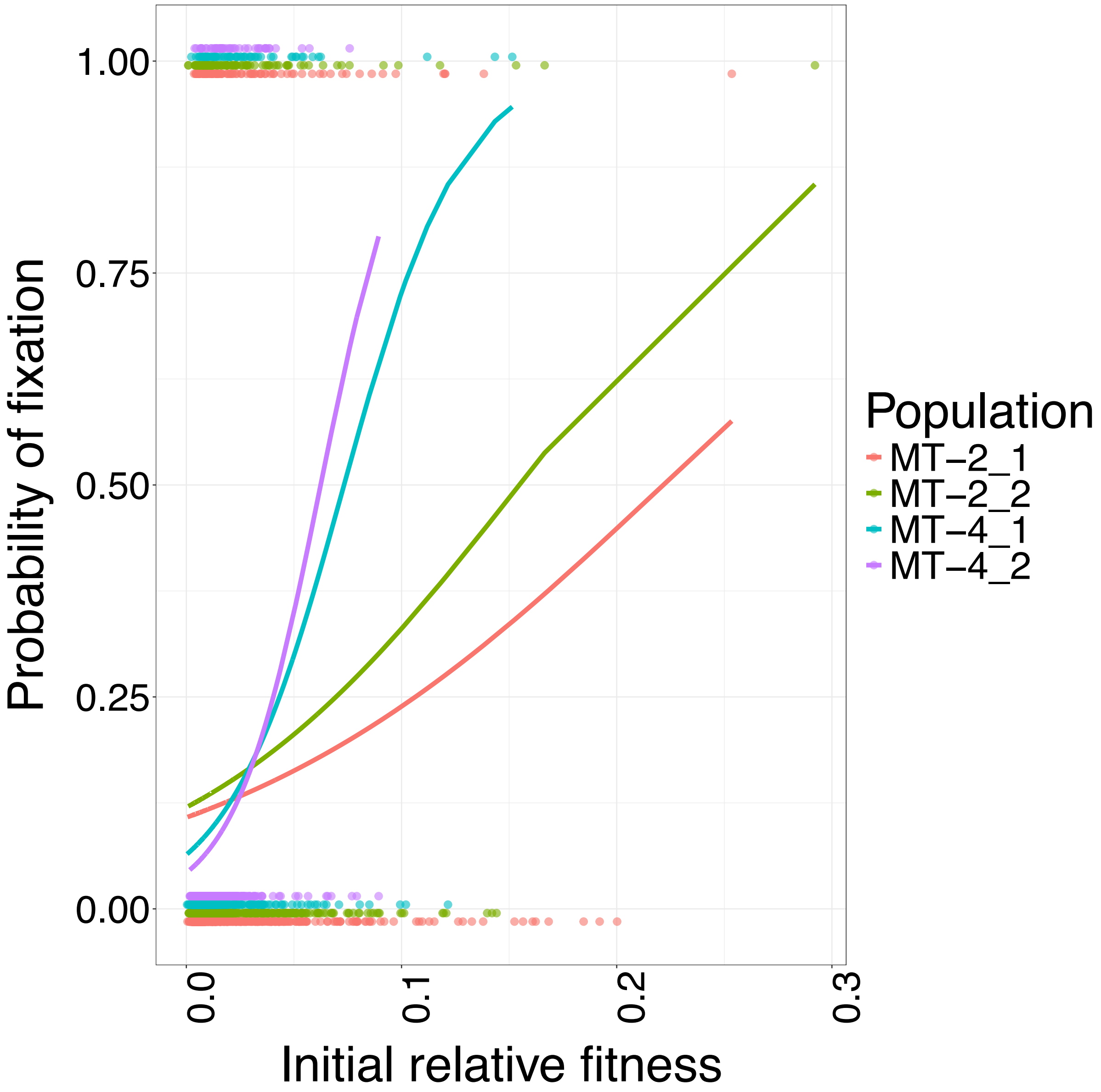


**Fig. S10. Probability of fixation based on initial relative fitness.** Mutations with larger initial relative fitness (i.e., larger initial increase in frequency in the population) were more likely to be fixed.

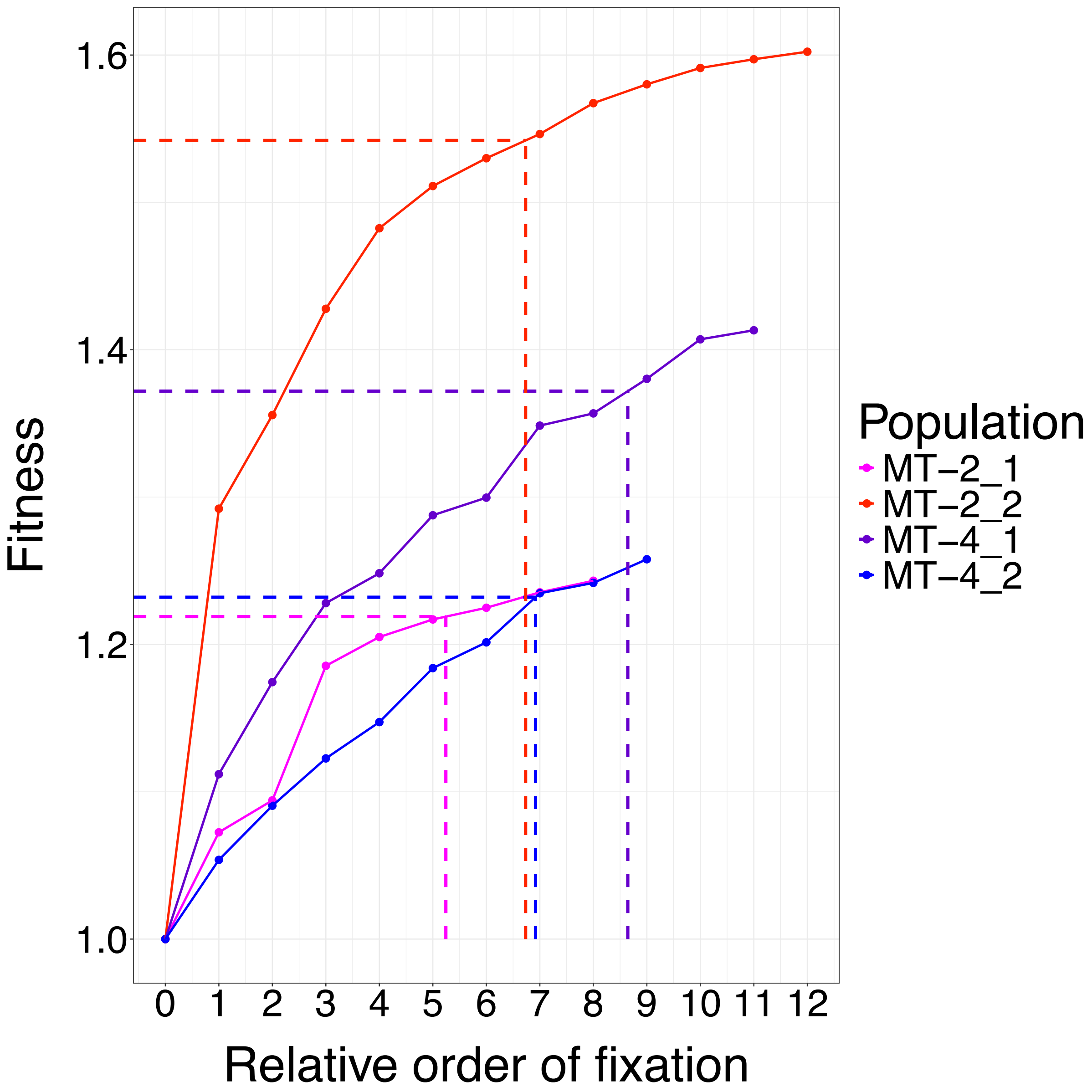


**Fig. S11. Cumulative fitness of evolving populations based on initial relative fitness of adaptive mutations.** The cumulative initial relative fitness conferred by each adaptive mutation in each lineage. The dashed lines depict the corresponding adaptive mutation where 90% of total fitness gain was achieved. The 90% of relative fitness gain was achieved by 6th (MT-2_1), 7th (MT-2_2), 7th (MT-4_2), and 9th (MT-4_1) adaptive mutation in each line. These mutations were fixed at transfers 360 (MT-2_1), 160 (MT-2_2), 290 (MT-4_2), and 350 (MT-4_1), respectively.

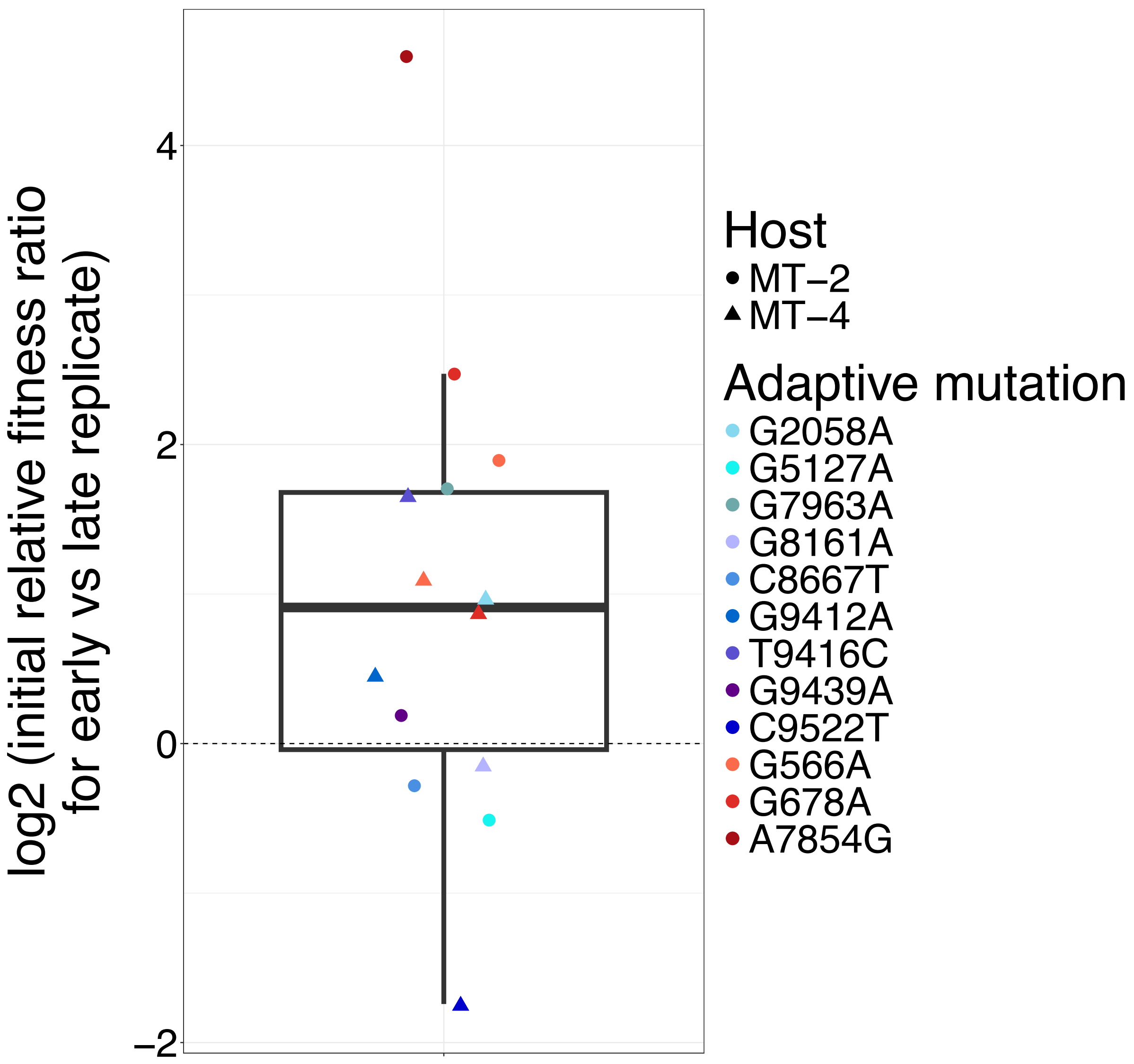


**Fig. S12. Ratio of initial relative fitness for adaptive mutations in early vs late replicate.** Adaptive mutations that occurred in both replicates of a given host and for which initial relative fitness data were available were identified. We then compared the ratio of initial relative fitness for the same adaptive mutation between the replicate in which it fixed earlier and the replicate in which it fixed later. Adaptive mutations that fixed earlier in a replicate had significantly higher initial relative fitness ($P < 0.05$, Wilcoxon rank-sum test).

### GLMER Model Summary for MT−2 and MT−4

*Negative Binomial Mixed Effects Models*

| Model | Term | Estimate | SE | z_value | p_value | Sig |
|---|---|---|---|---|---|---|
| MT−2 | (Intercept) | 2.7933 | 0.1431 | 19.520 | < 0.001 | *** |
| MT−2 | bin_mid_scaled | 0.4471 | 0.1798 | 2.487 | 0.01290 | * |
| MT−2 | groupHighly−parallel | −2.2165 | 0.3116 | −7.113 | < 0.001 | *** |
| MT−2 | bin_mid_scaled:groupHighly−parallel | −0.9994 | 0.3562 | −2.806 | 0.00502 | ** |
| MT−4 | (Intercept) | 1.4318 | 0.1662 | 8.615 | < 0.001 | *** |
| MT−4 | bin_mid_scaled | 0.6380 | 0.1769 | 3.607 | < 0.001 | *** |
| MT−4 | groupHighly−parallel | −0.7518 | 0.2898 | −2.594 | 0.00948 | ** |
| MT−4 | bin_mid_scaled:groupHighly−parallel | −1.1649 | 0.3115 | −3.740 | < 0.001 | *** |

| Model | AIC | BIC | logLik | Deviance | N_obs | N_groups | Theta | Marginal_R2 |
|---|---|---|---|---|---|---|---|---|
| MT−2 | 112.0 | 118.0 | −50.0 | 18.1 | 20 | 2 | 7.1608 | 0.874 |
| MT−4 | 86.1 | 92.1 | −37.0 | 22.6 | 20 | 2 | 106169.3779 | 0.638 |

*Note: Marginal $R^2$ quantifies the variance explained by fixed effects only.*

*Both models exhibit boundary/singular fits (random effect variance ... 0)*

**Fig. S13. Results and performance of generalized linear mixed-effects model.** Count of fixed mutations were modeled using negative binomial distribution based on time and occurrence group (line-specific and highly-parallel) as fixed terms and replicates as random effect. Two separate models were fit using the following formula for each host environment: $\text{count\_data} \sim \text{time} + \text{occurrence\_group} + \text{time} * \text{occurrence\_group} + (1|\text{replicate})$. As the interaction term between time and occurrence_group is significant in both models, we conclude that in both host environment the temporal patterns between line-specific and highly-parallel fixed mutations were the opposite. Line-specific fixed mutations increased by multiplicative factor of $e^{0.4471}$ and $e^{0.6380}$, while highly-parallel ones decreased by multiplicative factor of $e^{-0.5523}$ and $e^{-0.5269}$ per 100 transfer in MT-2 and MT-4, respectively.

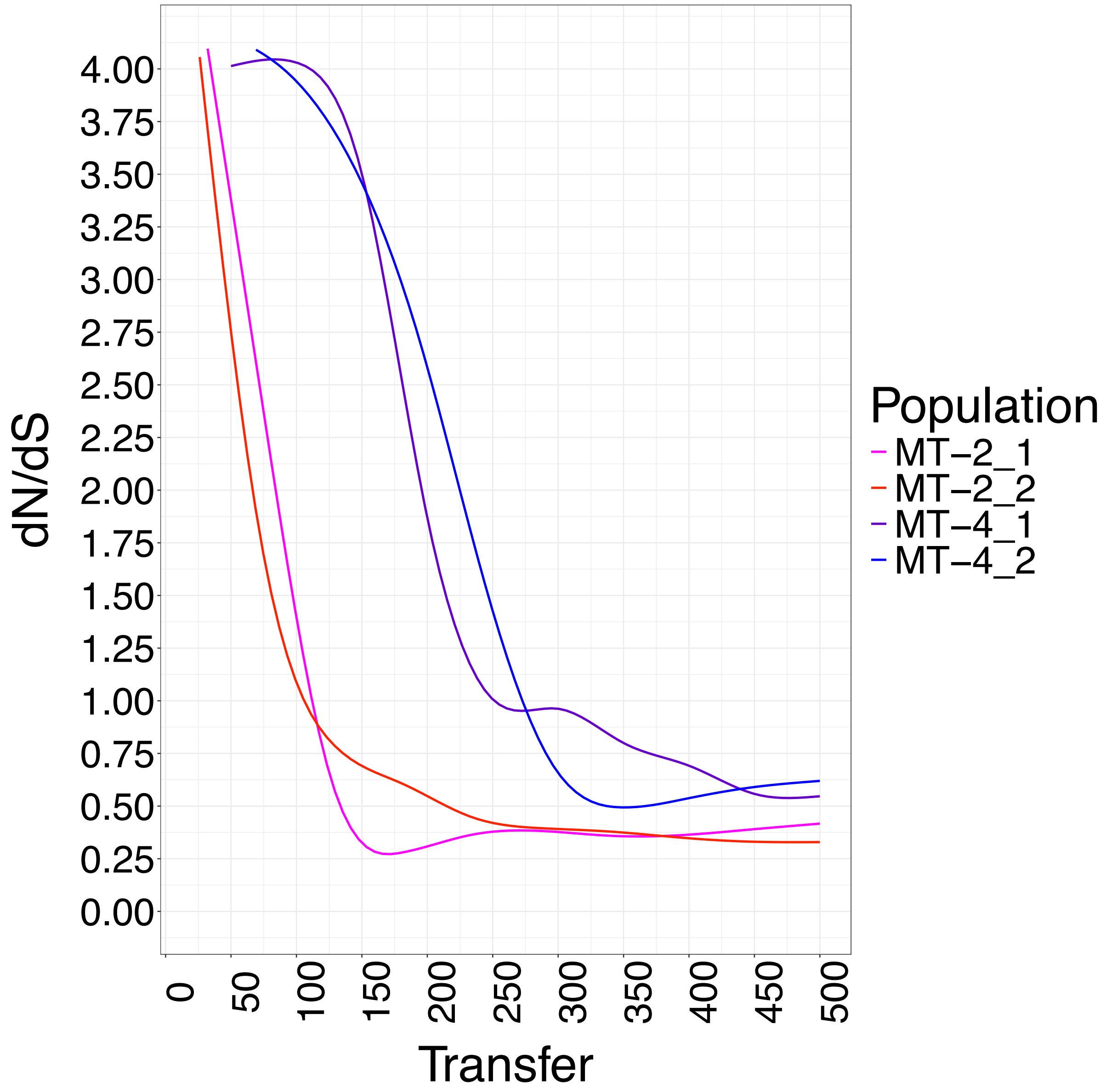


**Fig. S14. dN/dS ratio of fixed mutations overtime.** Early fixed mutations were predominantly nonsynonymous. The ratio between nonsynonymous and synonymous mutations per site changed in favor of the later by transfer 100 and transfer 250 in MT-2 and MT-4 replicates, respectively. The ratio seems to be stabilized at around 0.375 and 0.67 for MT-2 and MT-4 replicates, respectively.

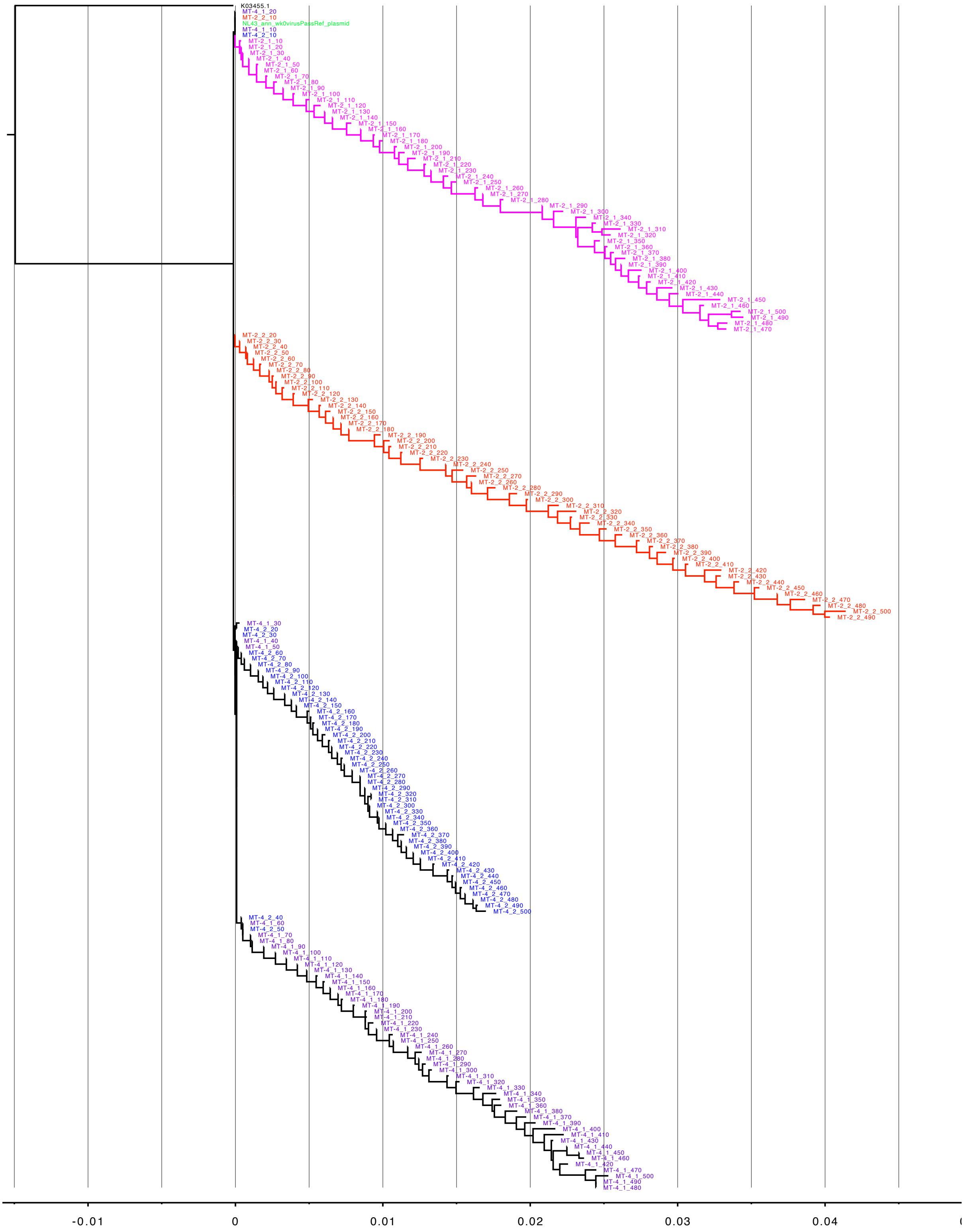


**Fig. S15. Reconstructed phylogeny.** The phylogenetic tree was reconstructed by maximum likelihood approach and based on consensus sequence of each sample population. The tree is mostly monophyletic, but rarlier samples were often misassigned to other clades due scarcity of mutations which were often shared between two lines. Overall, MT-2 replicates diverged at a faster rate. Even though the divergence rate was mostly constant over time, between transfers 260 and 310 the divergence rate was double the normal rate in MT-2_1. HXB2 sequence was used as an outgroup taxon.

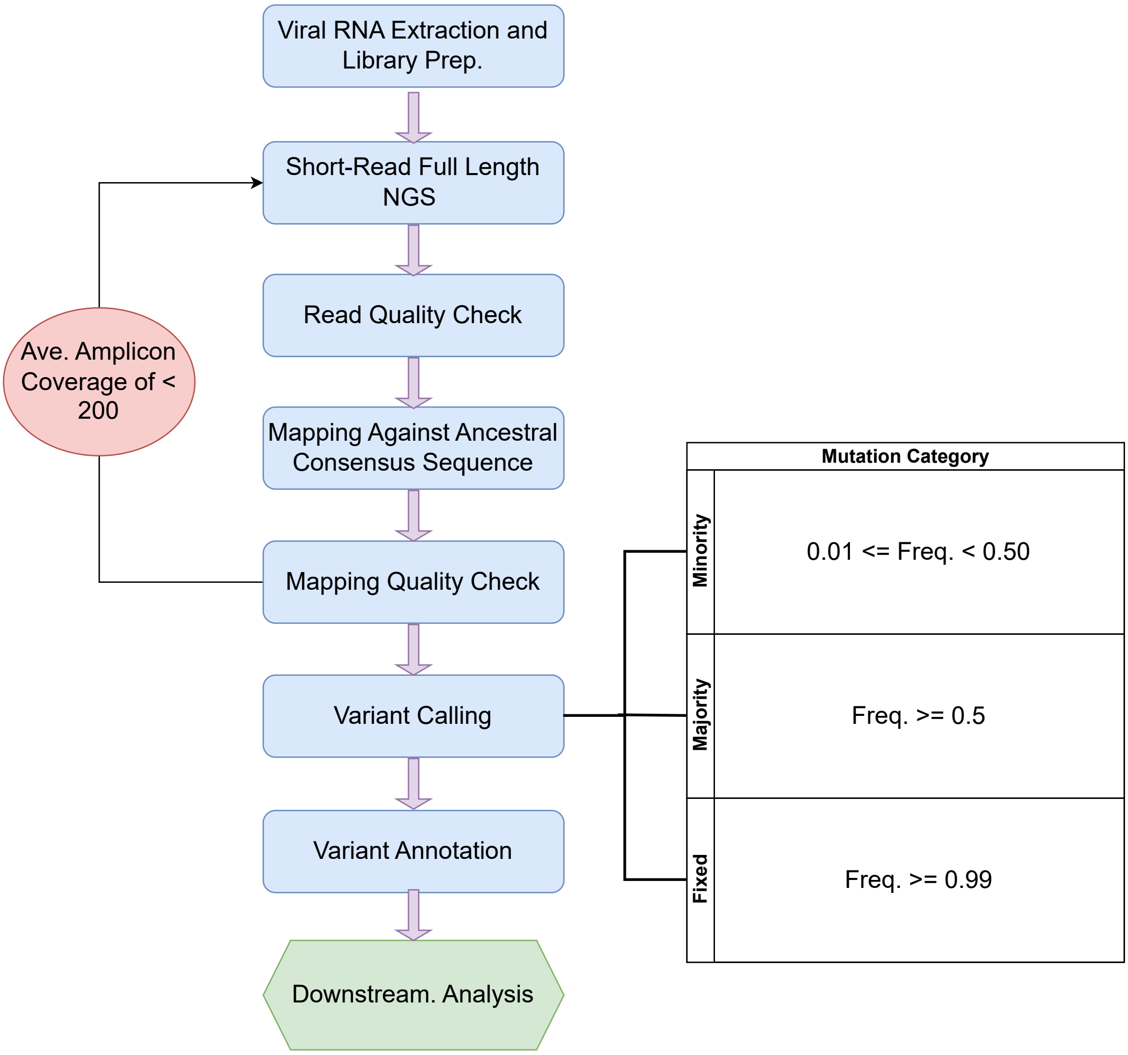


**Fig. S16. Sequencing workflow.** Schematic flowchart of the sequencing workflow. For details, please refer to Methods.

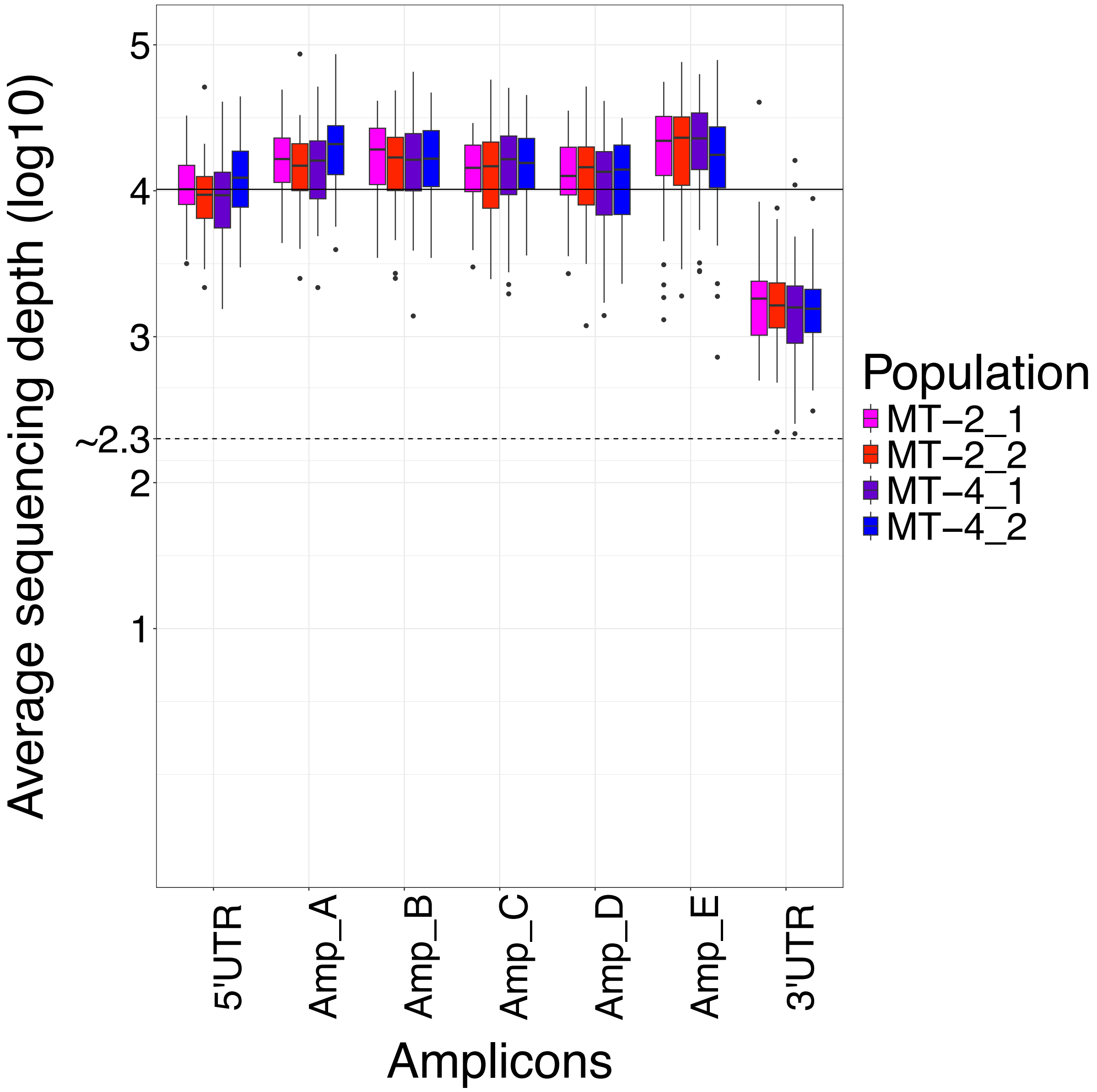


**Fig. S17. Sequence coverage per amplicon**. Five overlapping amplicons were designed to cover the whole length of HIV-1 genome. Average sequencing depth was measured per amplicon and per the two ends of the viral genome after mapping reads against the ancestral sequence. If the average sequencing depth of an amplicon for a sample was below 200 (the dashed line), the sequencing of the sample would be repeated. The average sequencing depth for any nucleotide in the genome was ~10,000 (the solid line).
